# Succinylation Links Metabolic Reductions to Amyloid and Tau Pathology

**DOI:** 10.1101/764837

**Authors:** Yun Yang, Victor Tapias, Diana Acosta, Hui Xu, Huanlian Chen, Ruchika Bhawal, Elizabeth Anderson, Elena Ivanova, Hening Lin, Botir T. Sagdullaev, William L. Klein, Kirsten L. Viola, Sam Gandy, Vahram Haroutunian, M. Flint Beal, David Eliezer, Sheng Zhang, Gary E. Gibson

**Author notes:** **Author Contributions** Y.Y. and G.G. conceived the research program and designed the experiments. V.H. contributed to the patient consent, collection of samples. V.H., X.H. and E.A. processed the brain samples. R.B. and S.Z. performed nanoLC-MS/MS analysis. Y.Y. performed data analyses. Y.Y. and X.H. performed biochemical experiments. Y.Y., X.H., and H.C. performed the cell experiments. X.H., H.C., E.I. and B.T.S. performed immunofluorescence on the rotenone treated cells and analyzed the data. D.A. and D.E. designed and performed NMR analysis, processed and interpreted the data and prepared figures. V.T. and M.F.B. participated in the design and conceptualization of the animal study, performed the experiments, analyzed the data, and prepared the figures. H.L. participated in the experimental design and write-up. S.G., W.K. and K.V. provided antibodies and contributed to the design. Y.Y. and G.G. wrote and edited the manuscript. All authors discussed the results, and Y.Y., G.G., S.Z., D.E., S.G., V.H., V.T., B.T.S, H.L. and E.I. contributed to the writing. All authors read and approved the manuscript.

## Abstract

Abnormalities in glucose metabolism and misfolded protein deposits composed of the amyloid-β peptide (Aβ) and tau are the three most common neuropathological hallmarks of Alzheimer’s disease (AD), but their relationship(s) to the disease process or to each other largely remains unclear. In this report, the first human brain quantitative lysine succinylome together with a global proteome analysis from controls and patients reveals that lysine succinylation contributes to these three key AD-related pathologies. Succinylation, a newly discovered protein post-translational modification (PTM), of multiple proteins, particularly mitochondrial proteins, declines with the progression of AD. In contrast, amyloid precursor protein (APP) and tau consistently exhibit the largest AD-related increases in succinylation, occurring at specific sites in AD brains but never in controls. Transgenic mouse studies demonstrate that succinylated APP and succinylated tau are detectable in the hippocampus concurrent with Aβ assemblies in the oligomer and insoluble fiber assembly states. Multiple biochemical approaches revealed that succinylation of APP alters APP processing so as to promote Aβ accumulation, while succinylation of tau promotes its aggregation and impairs its microtubule binding ability. Succinylation, therefore, is the first single PTM that can be added in parallel to multiple substrates, thereby promoting amyloidosis, tauopathy, and glucose hypometabolism. These data raise the possibility that, in order to show meaningful clinical benefit, any therapeutic and/or preventative measures destined for success must have an activity to either prevent or reverse the molecular pathologies attributable to excess succinylation.

## Introduction

Misfolded protein deposits of the amyloid beta peptide (Aβ)^1, 2^ and the microtubule-associated protein tau (tau)^3^ are central pathological features in Alzheimer’s Disease (AD), while reduced brain glucose metabolism and synaptic density are more highly correlated with the development of clinical cognitive dysfunction^4^. Preclinical research shows that diminished glucose metabolism exacerbates learning and memory deficits, concurrent with the accumulation of Aβ oligomers and plaques^5^, and misfolded, hyperphosphorylated tau^6, 7^. However, the interrelationships between and among these key pathological processes are largely unknown.

The decline in brain glucose metabolism in AD correlates with a reduction in the α-ketoglutarate dehydrogenase complex (KGDHC)^8^, a key control point in the tricarboxylic acid (TCA) cycle. In yeast^9^ and cultured neurons^10, 11^, reduction in KGDHC activity leads to a wide-spread reduction in regional brain post-translational lysine succinylation, a recently discovered post-translational modification (PTM). Studies of organisms deficient in NAD^+^-dependent desuccinylase sirtuin 5 (SIRT5)^12^ provide evidence of the regulatory importance of succinylation in metabolic processes^13–17^. However, the role of succinylation in metabolic pathways of the human nervous system or in neurodegenerative diseases is unknown. Our study represents the first to report the human brain succinylome and characterize its changes in AD. The results suggest that succinylation links the AD-related metabolic deficits to structural, functional and pathological changes in APP and tau.

### Succinylome and proteome of human brain

Analysis of two cohorts each consisting of brain tissues from five controls and five AD patients (patient information is provided in **Supplementary Table 1**) was performed in order to maximize our chances of optimizing the precision and reproducibility of the determinations of the succinylome (**Figure 1a, b**) and the proteome (**Figure 1c, d**). When the two independent cohorts were taken together, 1,908 succinylated peptides from 314 unique proteins were identified across a total sample size of 20 brains (**Figure 1b**). The parallel global proteomic analysis detected 4,678 proteins (**Figure 1d**). Nearly all of the succinylated proteins identified during the study were found in the global proteome of the same samples (**Figure 1e**).

**Figure 1.**
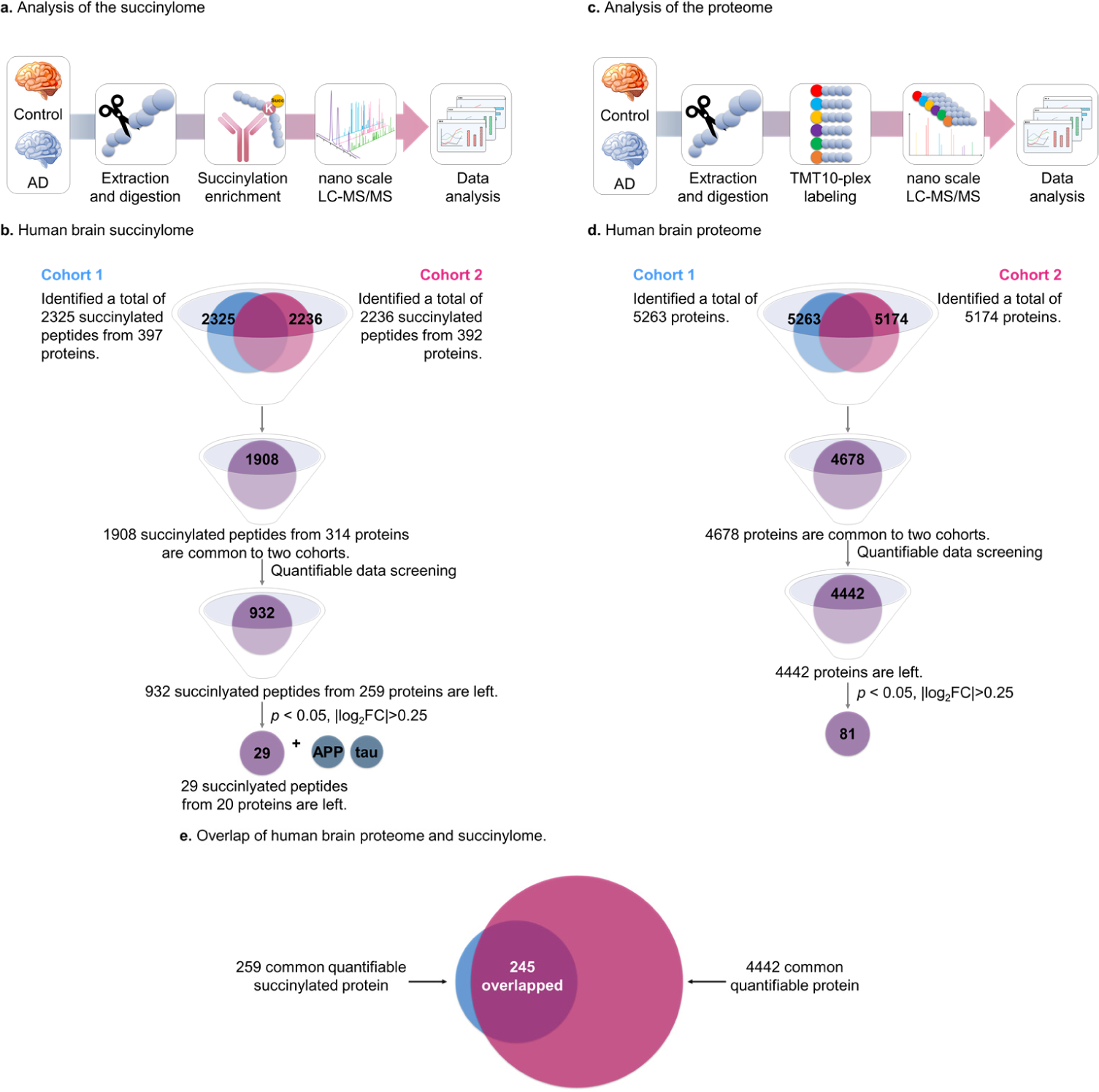
Global analysis of protein lysine succinylation and proteomic profiles in human brains.

a. A schematic diagram of the workflow for investigation of human brain lysine succinylome by label-free quantitation (See methods section).
b. After quantitative data screening and mining, the combined results from 20 brain samples in two batches revealed 932 common succinylated peptides quantified from 259 proteins (**Supplementary Table 4**).
c. A schematic diagram of the workflow for quantitative proteomics of human brain by Tandem mass tags (TMT) labeling analysis (See methods section).
d. After quantitative data screening and mining, the combined results from 20 brain samples in two batches revealed 4,442 common proteins in both AD and controls (**Supplementary Table 6**). Eighty-one proteins showed significant alterations between samples patients with AD and controls.
e. The overlap between succinylomes and proteomes. Nearly all of the succinylated proteins were also identified in its global proteomic analysis.

Subcellular localization analysis of the 314 succinylated proteins from 20 human brains facilitates an understanding of the implications of succinylation for cell function (**Figure 2a** and **Supplementary Table 2**). Succinylated proteins were one-to-many mapped to multiple subcellular compartments. Among those, mitochondrial proteins were the most heavily succinylated (**Figure 2b**). About 73% (229/314) of the succinylated proteins were mitochondrial. The pyruvate dehydrogenase complex (PDHC) E1 component subunit alpha (PDHA1), which links glycolysis to the TCA cycle, was succinylated extensively. All eight enzymes of the TCA cycle in the mitochondrial matrix and their multiple subunits, were also succinylated extensively. Succinylated proteins were also associated with the cytosol (30%, 95 proteins) and nucleus (23%, 73 proteins) (**Figure 2b**). The overall distribution resembled that reported for succinylated proteins in mouse liver^16, 17^.

**Figure 2.**
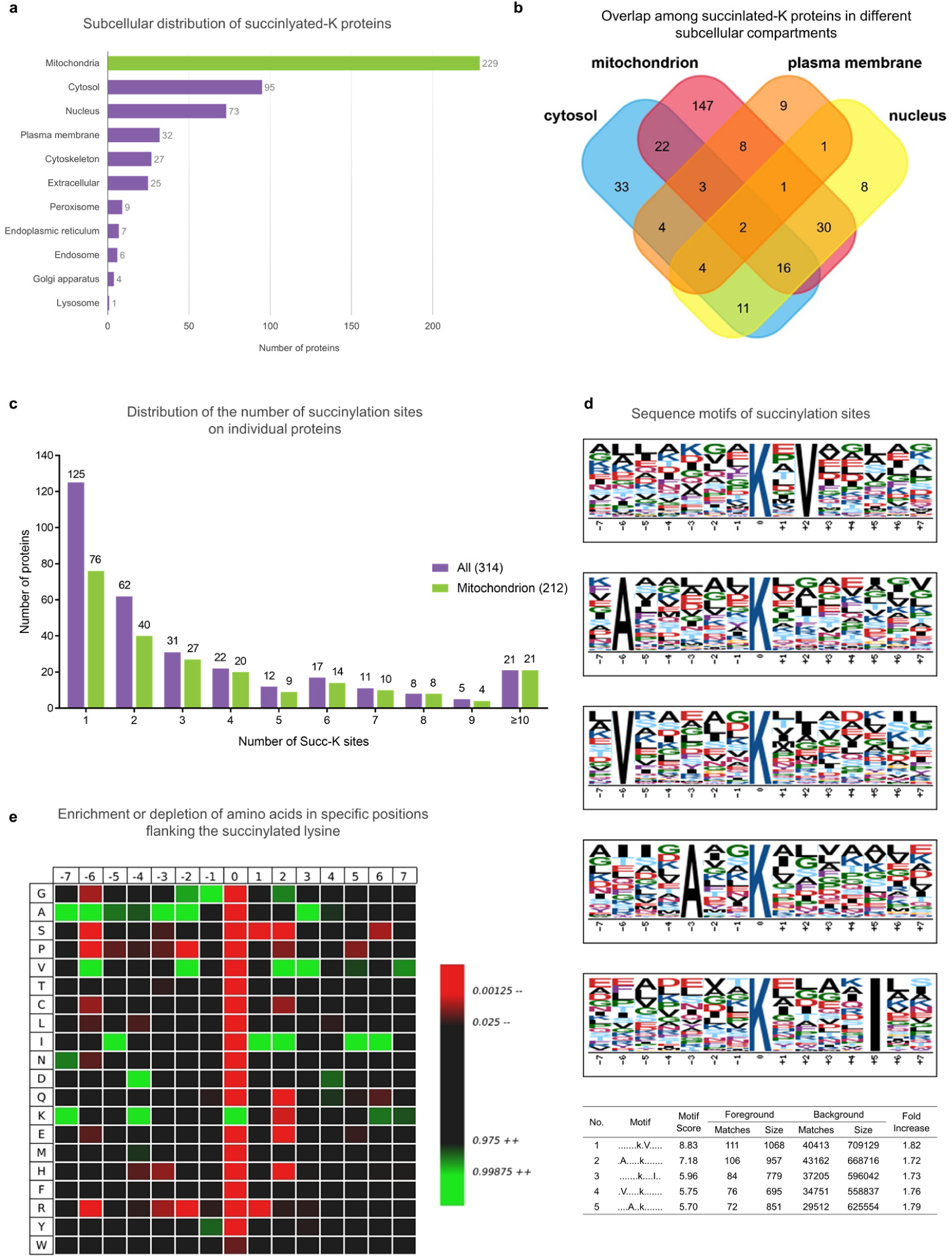
Subcellular distribution of lysine succinylation proteins in human brains.

a. Subcellular distribution of succinlyated-K proteins identified by Cytoscape and stringAPP software. The majority of succinylated-K proteins are mitochondrial.
b. Overlap of succinlated-K proteins located in the mitochondrion, nucleus, cytosol and plasma membrane. The details of the subcellular distribution of individual proteins are shown in **Supplementary Table 2**.
c. The extent of succinylation of individual proteins and their enrichment in mitochondria. Distribution of the number of succinylation sites per protein in all of the succinylated proteins (purple bars) or succinylated mitochondrial proteins (green bars) as classified by Cytoscape and stringAPP.
d. The succinylation sites were analyzed for seven amino acids up- and down-stream of the lysine residue using Motif-X. The height of each letter corresponds to the frequency of that amino acid residue in that position. The central blue K refers to the succinylated lysine.
e. Heat map of the 15 amino acid compositions of the succinylated site showing the frequency of the different amino acids in specific positions flanking the succinylated lysine. The different colors of blocks represent the preference of each residue in the position of a 15 amino acid-long sequence context (green indicates greater possibility, while red refers to less possibility).

The number of succinylation sites per protein varied from 1 to 23 (**Figure 2c** and **Supplement Table 2**), with 40% (125/314) having one succinylated site, 20% (60/314) having two, and the remaining 40% (127/314) having three or more. Eighty-nine percent of proteins with more than two succinylated lysines were mitochondrial. Moreover, the most extensively succinylated proteins with over ten distinct succinylated sites/peptides were all mitochondrial proteins, and 61% (14/21) of these are exclusively mitochondrial proteins including isocitrate dehydrogenase (IDH2), fumarate hydratase (FH) and malate dehydrogenase (MDH2) (see **Supplement Table 2** in red). In general, these succinylated proteins typically appeared in metabolism-associated processes and were linked to multiple disease pathways in KEGG enrichment analysis (**Extended Data Figure 1** and **Supplement Table 3**).

Since no specific motifs for lysine succinylation in human cells have been reported, a succinylation motif analysis of all 1908 succinylated peptides using Motif-X^18^ was used to assess whether specific motif sites exist. A total of five conserved motifs were identified (**Figure 2d**). A survey of these motifs suggested that non-polar, aliphatic residues including alanine, valine and isoleucine surround the succinylated lysines. Succinylated lysine site analysis revealed a strong bias for alanine residues, which is consistent with motifs identified in tomato^14^. IceLogo^19^ heat maps assessed the preference of each residue in the position of a 15 amino acid-long sequence context (**Figure 2e**). Isoleucine was detected downstream of lysine-succinylation sites, while alanine and lysine, two of the most conserved amino acid residues, were found upstream. Meanwhile, valine residues occurred upstream and downstream. By contrast, there was only a very small chance that tryptophan, proline or serine residues occurred in the succinylated peptides.

### Succinylome and proteome changes in AD brains

Completion of the human brain succinylome and global proteome analyses allowed direct comparison between brains form controls and AD patients. Of 1,908 succinylated peptides identified in two independent analyses (n = 5 control brains; n = 5 AD brains), 932 succinylated peptides were quantifiable (**Figure 1a**). A volcano graph analysis revealed that the succinylation of 434 unique peptides declined with AD while the abundance of 498 unique succinylated peptides was increased (**Figure 3a** and **Supplement Table 4**). Succinylation of 29 peptides (from 20 proteins) differed significantly (two-tailed Student’s t-test, *p* < 0.05) between AD and controls (**Figure 3a, b**). Succinylation of ten peptides increased with AD while succinylation of 19 peptides decreased.

**Figure 3.**
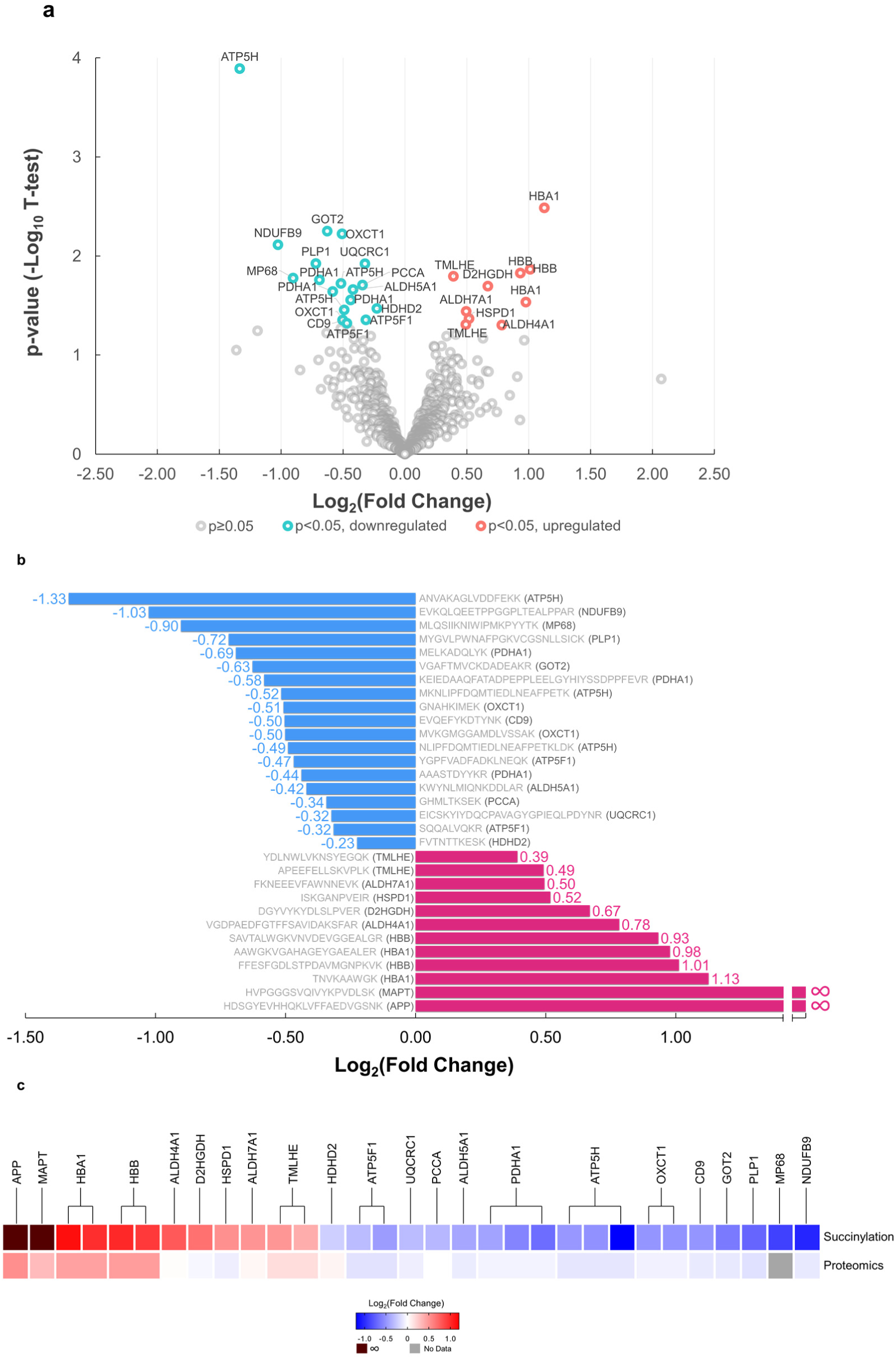
Comparison of the succinylome of brains from ten controls and ten patients with AD reveal many specific differences (*p*<0.05, two-sided Student’s t-test).

a. Volcano plot of 932 brain protein peptide succinylation in controls and AD patients. The signal detection result shows the magnitude (log_2_Fold Change, x-axis) and significance (− log_10_ *p-*value, y-axis) for brain succinylation changes associated with AD. Each spot represents a specific succinylated peptide. Green symbols to the left of zero indicate succinylated peptides that are decreased significantly while red symbols to the right of zero indicate succinylated peptides that are upregulated significantly in AD brains (*p*<0.05, two-sided Student’s t-test).
b. Peptides with significant differences in succinylation between control and AD brains. Decreases (blue bars) or increases (red bars) from the control succinylome are depicted as relative fold change. The sequence of the peptide and the name of the gene to which the peptides belong is noted for each bar.
c. Comparison of the AD-related changes in global proteome and succinylome. The succinylated peptides from the succinylome were clustered based on their proteins. For each protein, its relative fold change in succinylome and global proteome of AD cases versus controls is shown.

Proteomic analysis of 20 samples in two cohorts (**Figure 1c)** showed that of the 4,678 identified proteins, 4,442 common proteins were quantifiable in both AD and controls (**Figure 1d** and **Extended Data Figure 2a, b**). Comparison of the succinylome with the proteome demonstrated that the AD-related changes in succinylation of these peptides were only weakly correlated with -- and therefore unlikely to be due to -- changes in corresponding protein levels (**Figure 3c**). The proteomic analysis revealed that 81 proteins changed significantly (two-tailed Student’s t-test, *p* < 0.05 and |log_2_FC| > 0.25). Eight proteins decreased in brains from AD patients, while 73 proteins increased (**Extended Data Figure 2a**).

The overwhelming majority (16/19) of the peptides with AD-related decreases in succinylation were mitochondrial, and more than half of them showed exclusive localization in mitochondria (**Supplementary Table 5**). A novel association of the ATP5H/KCTD2 locus with AD has been reported^20^, and ATP-synthase activity declines in AD brains^21^. In line with these findings, we identified the maximal AD-related decrease (−1.33 in log2FC) in ATP synthase subunit d (ATP5H), with two additional peptides from ATP5H down at −0.52 and −0.49 in log2FC. Moreover, two peptides from another subunit, namely ATP synthase subunit b (ATP5F1), also decreased (log2FC at −0.47- and −0.32) in AD brains. Succinylation of three lysine residues (Lys77, Lys244 and Lys^344^) of PDHA1 also decreased significantly with AD (**Figures 3a, 3b**).

The largest AD-related increases in succinylation were in non-mitochondrial proteins (**Figures 3a, 3b**). Succinylation of four peptides from brain cytosolic and/or extracellular hemoglobin subunits alpha and beta increased by 1.91- (0.978 in log2FC) to 2.18-fold (1.127 in log2FC) with AD. Strikingly, two extra-mitochondrial peptides with the largest AD-related increases in succinylation were from two proteins critical to AD pathology: APP and tau. Both proteins were highly succinylated at critical sites in nine out of ten AD brain samples, but no succinylation of APP or tau was detectable in any control brains (**Figures 5, 6**).

### Subcellular responses of succinylation to impaired mitochondrial function

Subcellular succinylation in response to perturbed mitochondrial function was determined by compromising mitochondrial function of HEK293T cells by mild inhibition of complex I (20-minute-treatment) followed by determining the effects on succinylation. Impaired mitochondrial function diminished general succinylation in whole cell lysates and mitochondrial fractions (**Figure 4a**), consistent with previous findings in N2a cells^11^. However, mitochondrial dysfunction increased succinylation of 30-70 kDa proteins in the non-mitochondrial fractions. We previously demonstrated that mitochondrial dysfunction can alter mitochondrial/cytosolic protein signaling^22^. Here we extend this line of investigation by showing that mitochondrial dysfunction resulted in a release of mitochondrial proteins including all subunits of PDHC and KGDHC (**Figure 4b, c**). This was not due to disruption of the mitochondrial integrity because cytochrome c oxidase subunit 4 isoform 1 (CoxIV), an integral membrane protein in mitochondria, did not increase in the cytosol fraction. Confocal microscopy further confirmed that rotenone caused a redistribution of mitochondrial proteins without mitochondrial lysis, as mitochondria were clearly outlined by CoxIV immunolabeling. Rotenone treatment increased the amount of the cytosolic E2k component of KGDHC (DLST) outside of mitochondia defined by CoxIV (**Figure 4d**). Thus, impaired mitochondrial function induced a metabolic disturbance leading to an increased leakge of mitochondrial proteins into the cytosol, including DLST. DLST, being a succinytransferase^10^ and a succinyl-CoA generator^23^, increased succinylation in non-mitochondrial fractions.

**Figure 4.**
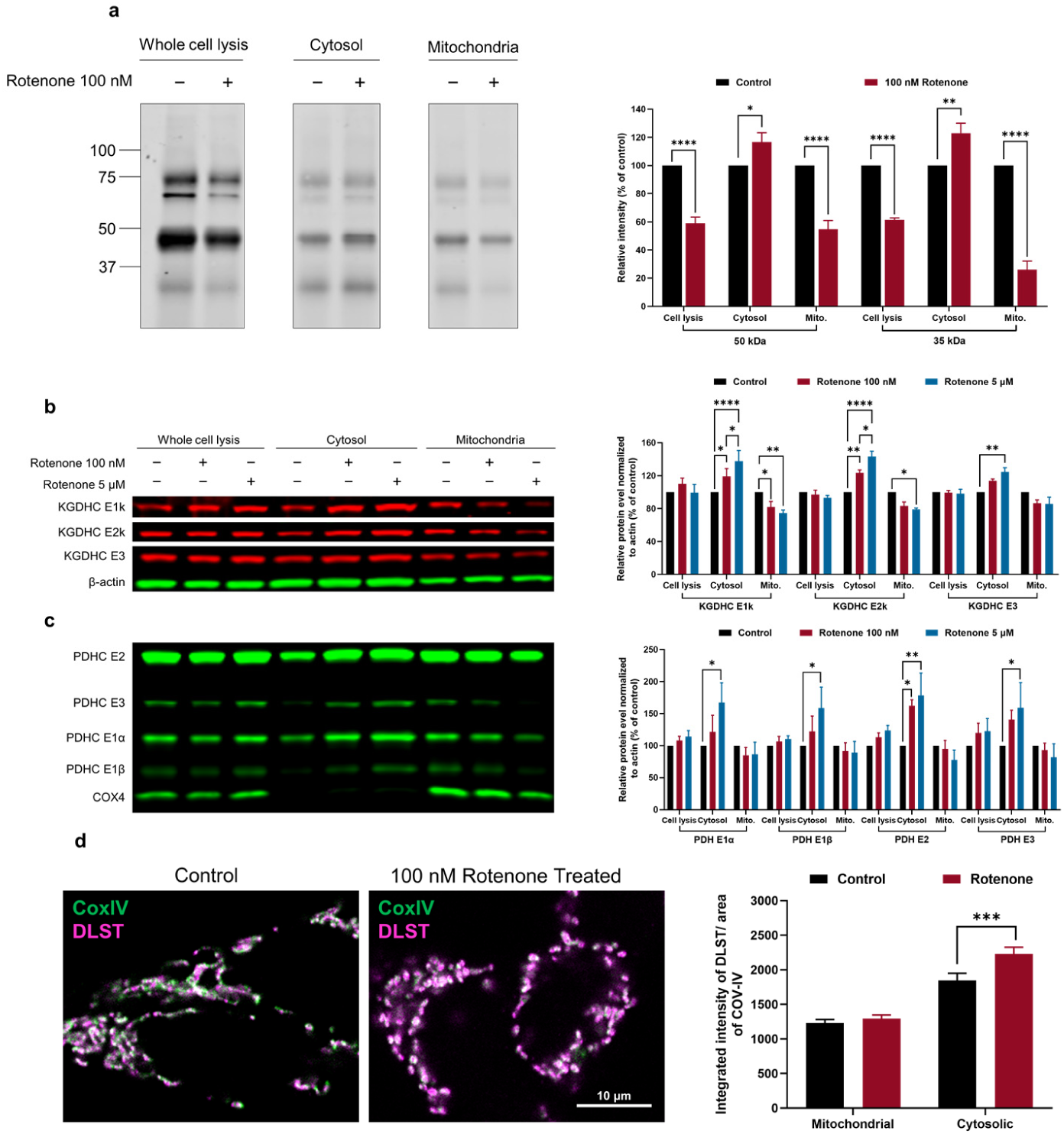
Impairing mitochondrial function altered succinylation and protein distribution in the whole cell as well as in the mitochondria and non-mitochondrial fractions.

a. The effects of rotenone (100 nM/20 min) on succinylation in HEK293T cells. After separation, mitochondrial and non-mitochondrial fractions were immune-precipitated with anti-succinyllysine antibody and separated by SDS-PAGE followed by Western blotting. The data from three different replicate experiments were expressed as the mean with error bars from standard error of the mean (SEM) (n = 3, ****: *p* < 0.0001, **: *p* < 0.01, *: *p* < 0.05, two-way ANOVA followed by Bonferroni’s multiple comparisons test).
b. The effects of rotenone (100 nM, 5 μM/20 min) on the distribution of KGDHC protein between mitochondria and non-mitochondrial fractions. The data from three different replicate experiments were expressed as the mean with error bars from SEM (n = 3, ****: *p* < 0.0001, **: *p* < 0.01, *: *p* < 0.05, two-way ANOVA followed by Tukey’s multiple comparisons test).
c. The effects of rotenone (100 nM, 5 μM/20 min) on the distribution of PDHC protein between mitochondria and non-mitochondrial fractions. The data from three different replicate experiments were expressed as the mean with error bars from SEM (n = 3, **: *p* < 0.01, *: *p* < 0.05, two-way ANOVA followed by Tukey’s multiple comparisons test).
d. Confocal microscope analysis results of DLST and mitochondrial mass. co-localization in HEK293T cells in response to the mitochondrial dysfunction. Magenta: DLST; Green: CoxIV; Error bars represent SEM deviation from the mean (n = 98 fields from 19 dishes, ***: *p* < 0.001, Tukey’s multiple comparisons test).

**Figure 5.**
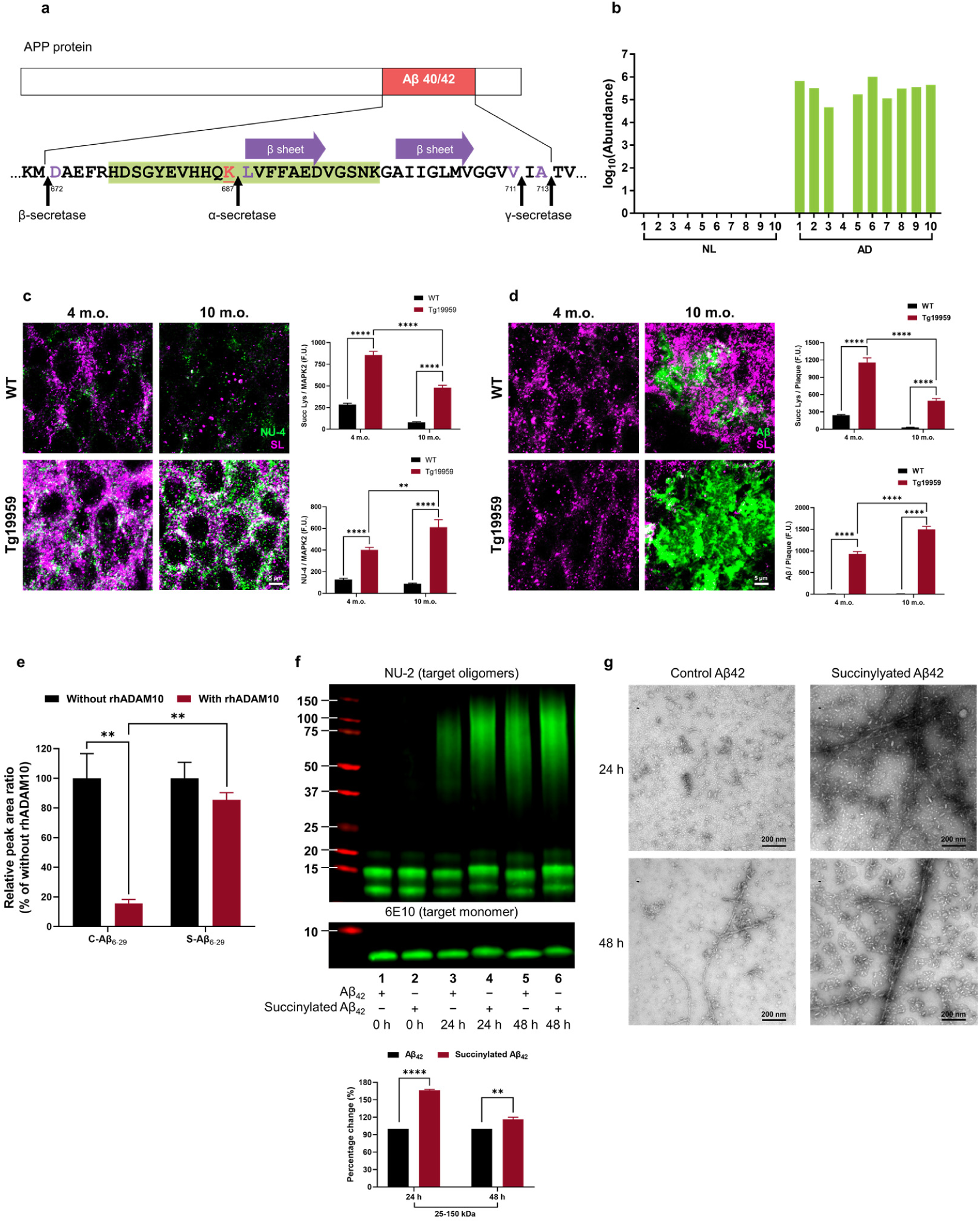
Succinylation occurs uniquely on APP from AD patients, in early stages of plaque formation in mouse models and disrupts APP processing.

a. Location and identity of succinylation K687 near the Aβ region. Residues are numbered according to APP770 sequence. Purple amino acids refer to α- or β- or γ-cleavage sites. The red underlined lysine refers to succinylated K687. Purple arrow represents the two central strands of the β-sheet (Leu688-Asp694 and Ala701-Val707). Green highlights the peptide identified in the MS.
b. Abundance of succinylation K687 found in brains from 10 controls and 10 AD patients. Data transformed by log_10_ (abundance) for normalization purposes and to facilitate presentation.
c. Confocal microscope analysis of the co-localization of succinylation and amyloid oligomers in the hippocampal CA1 region sections from 4-month-old and 10-month-old Tg19959 or WT mice. NU-4 (green) staining Aβ oligomers; pan-succinyl-lysine (magenta). Four mice per group. Data were expressed as the mean with SEM representative of the average of ∼900-1,000 MAP2 neurons or 60 Aβ plaques comprised in 3-4 different hippocampal sections per animal. The fluorescence intensity of succinyl lysine was normalized to the number of pyramidal neurons (****: *p* < 0.0001, **: *p* < 0.01, two-way ANOVA followed by Tukey’s multiple comparisons test).
d. Confocal microscope analysis of the co-localization of succinylation and plaque pathology in the hippocampal CA1 region sections from 4-month-old and 10-month-old Tg19959 or WT mice. Aβ (green) staining plaque; pan-succinyl-lysine (magenta). Four mice per group. Data were expressed as the mean with SEM representative of the average of ∼900-1,000 MAP2 neurons or 60 Aβ plaques comprised in 3-4 different hippocampal sections per animal. The fluorescence intensity of succinyl lysine was normalized to the number of pyramidal neurons (****: *p* < 0.0001, two-way ANOVA followed by Tukey’s multiple comparisons test).
e. Succinylation blocks α-cleavage. Peptides were incubated for 24 hrs with or without rhADAM10. Peak area ratio values were calculated and are shown relative to corresponding controls without rhADAM10. Each sample was run in triplicate and data were expressed as the mean with SEM (**: *p* < 0.01, two-way ANOVA followed by Bonferroni’s multiple comparisons test; except for one sample from the group of succinylated peptide without rhADAM10 was damaged).
f. Western blot analysis of succinylated and control Aβ_42_ from aggregation assay showed that the succinylation generates more oligomerized Aβ even after a long incubation. The data from two different replicate experiments were expressed as the mean with error bars from SEM (****: *p* < 0.0001, **: *p* < 0.01, two-way ANOVA followed by Bonferroni’s multiple comparisons test).
g. Two time points from aggregation assay were analyzed by negative-staining electron microscopy.

### Functional significance of succinylation of APP

AD-associated succinylation of APP occurred at a critical site (K687) in nine of ten brains from AD patients but not in controls (**Figure 5a, b**), and the following experiments demonstrated it to be pathologically important. In Tg19959 mice bearing human APP with two AD-related mutations, the early amyloid pathological changes appeared at 4 months (**Figure 5c** and **Extended Data Figure 3a**), and amyloid deposits developed by 10 months (**Figure 5d** and **Extended Data Figure 3b**). Double immunofluorescence staining with antibodies to pan-lysine-succinylation and to Aβ oligomers (NU-4)^24^ or to Aβ plaque (β-Amyloid (D3D2N)) revealed a very early increase in succinylation that appeared to paralleled oligomer formation and subsequent plaque formation in the hippocampus. These findings suggest that the APP succinylation might be involved in Aβ oligomerization and plaque formation throughout the development of plaque pathology *in vivo*.

In subsequent experiments, we tested the relationship between succinylation and APP processing by the secretase enzymes. K687-L688 is the APP α-secretase cleavage bond, and a missense mutation at K687N produces an early onset dementia^25^. Furthermore, global proteomics showed an increase of β-secretase (BACE1) abundance of 31% in AD brains compared to controls (**Supplementary Data Table 6**), while no changes occurred for either α-secretase or the SIRT family (**Extended Data Figure 2c**). Thus, succinylation of APP at K687 in AD may promote Aβ production by inhibiting α-secretase cleavage. To test this, synthetic peptides comprised of residues 6-29 in Aβ_42_ (numbering with respect to the N terminus of Aβ_42_), which span the α-secretase cleavage site, with or without succinylation at K16 (corresponding to K678 in APP), were assayed for α-secretase cleavage susceptibility. Recombinant human ADAM10 (rhADAM10) cleaved the native (control) peptide (substrate) with 84% efficiency, whereas no cleavage of its succinylated counterpart was detectable following a 24-hrs incubation (**Figure 5e**). Measurement of the two fragments that are produced by α-secretase activity confirmed a strong inhibition of α-secretase activity (**Extended data Figure 3c-g**).

Residue K16 (K687 in APP) is critical for both aggregation and toxicity of Aβ_42_^2,26^. Aβ oligomers are widely regarded as the most toxic and pathogenic form of Aβ^27^. To assess whether succinylation can directly alter Aβ oligomerization, aggregation of succinylated and non-succinylated Aβ_42_ was determined by anti-Aβ oligomer antibody NU-2^24^ and electron microscopy (EM). After 24 and 48 hrs incubation, succinylation promoted more robust Aβ oligomerization (**Figure 5f**). Moreover, the EM micrographs clearly revealed elevated levels of oligomeric, protofibrillar, and fibrillar Aβ^28^ in the succinylation group at t = 24 or 48 hrs (**Figure 5g**). These data revealed that succinylation of K687 of APP was a key molecular pathological underpinning that promoted Aβ oligomerization. Taken together, the accumulated data strongly suggest that succinylation of K678 might lead to an early-onset enhanced generation, oligomerization and plaque biogenesis, consistent with the effects of known genetic disease mutations at this site^25, 29^.

### Functional significance of succinylation of tau

Tau has two important nucleating sequences that initiate the aggregation process: PHF6 (residues 306-311) and PHF6* (residues 275-280) (**Figure 6a**)^30, 31^. PHF6* is located at the beginning of the second repeat (R2) and is only present in all four-repeat tau isoforms, while PHF6 is located at the beginning of the third repeat (R3) and is present in all tau isoforms. Tau succinylation on K311 within the PHF6 hexapeptide ^306^VQIVYK^311^ was detected in nine of ten AD brain samples but was undetectable in all control (**Figure 6b**). Acetylation of K280 of PHF6* in tau is a well-characterized^32^ modification that affects tau function^3^, and has become a prognostic factor and a new potential therapeutic target for treating tauopathies. Removal of residue K311 in PHF6 abrogated fibril formation^33^, but the structural and functional implications of K311 succinylation are unknown. Thus, exploring the influence of tau succinylation on K311 may be important as we seek to develop a comprehensive understanding of the effects.

**Figure 6.**
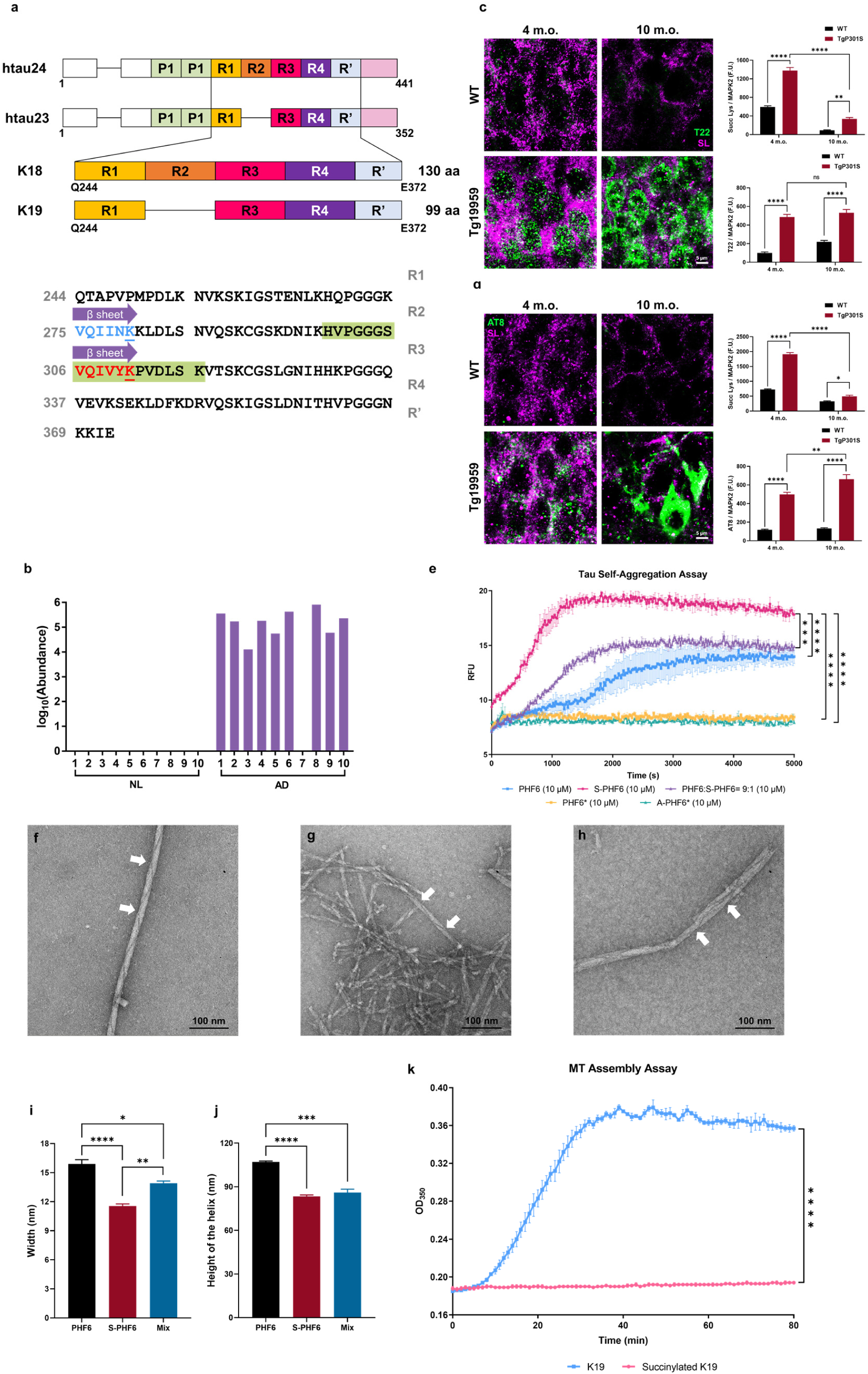
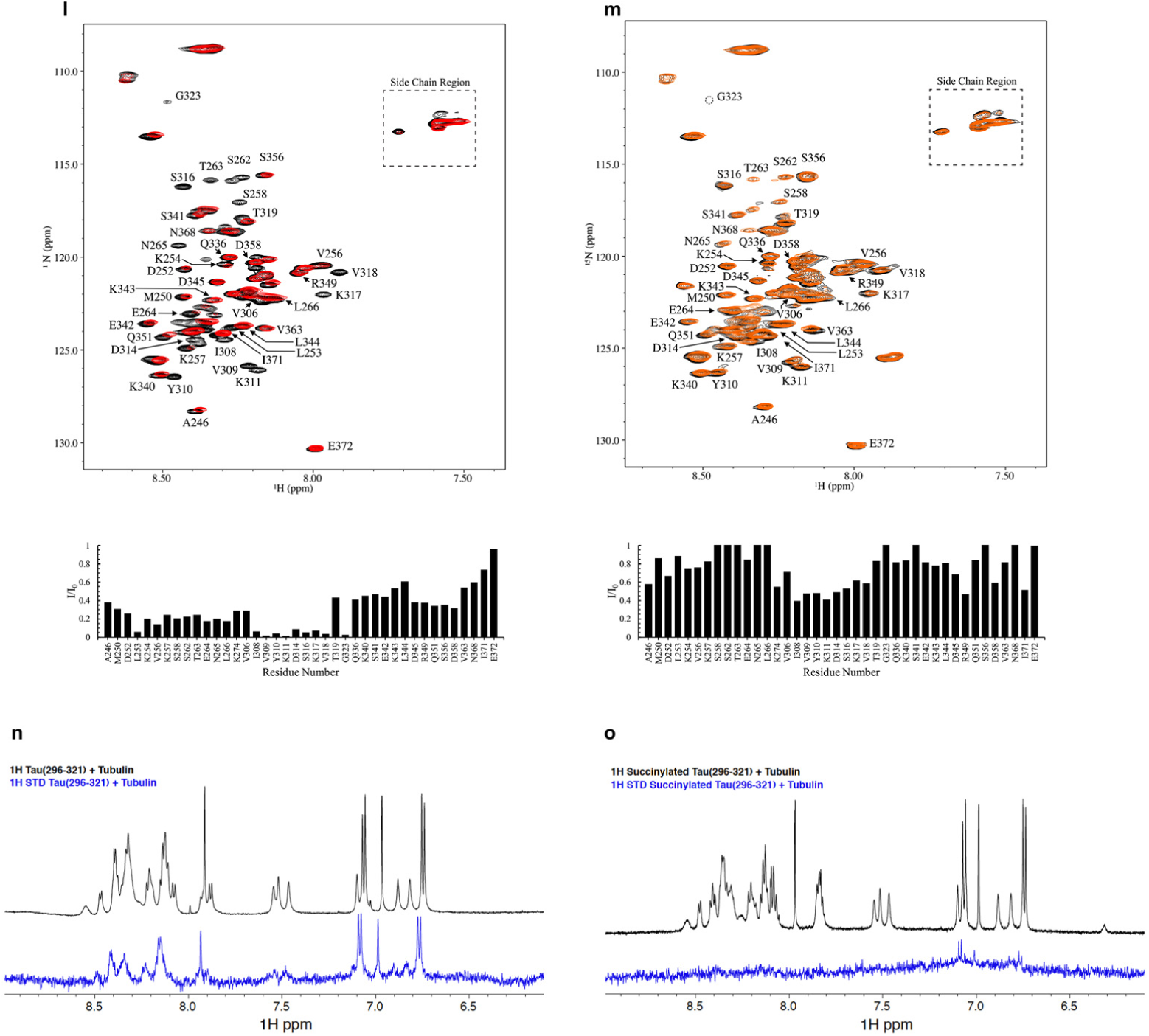
The unique succinylation of K311 on tau in brains from patients with AD promotes AD like features in tau pathology.

**a.** Domain structure of tau and the location of succinylation K311. The diagram shows the domain structure of htau23 and 24, which contain three and four repeats, respectively. The constructs K18 and K19 comprise four repeats and three repeats, respectively. Residues are numbered according to tau441 sequence. Purple arrow represents the two central strands of the β-sheet (PHF6*: Val275-Lys280, highlighted in blue, the blue underlined lysine refers to acetylated K280; PHF6: Val306-Lys311, highlighted in red, the red underlined lysine refers to succinylated K311). Green highlights the peptide identified by MS.
**b.** Abundance of succinylation K311 found in brains from ten controls and ten patients with AD. Data transformed by log_10_ (abundance) for normalization purposes and to facilitate presentation.
**c.** High confocal microscope analysis results of the co-localization of succinylation and tau oligomers in the hippocampal CA1 region sections from 4-month-old and 10-month-old TgP301S or WT mice. T22 (green) staining tau oligomers; pan-succinyl-lysine (magenta). Four mice per group. Data were expressed as the mean with SEM representative of the average of ∼900-1000 MAP2 neurons or 60 Aβ plaques comprised in 3-4 different hippocampal sections per animal. The fluorescence intensity of succinyl lysine was normalized to the number of pyramidal neurons (****: *p* < 0.0001, **: *p* < 0.01, two-way ANOVA followed by Tukey’s multiple comparisons test).
**d.** High confocal microscope analysis results of the co-localization of succinylation and phospho-tangle pathology in the hippocampal CA1 region sections from 4-month-old and 10-month-old TgP301S or WT mice. AT8 (green) staining phospho-tau; pan-succinyl-lysine (magenta). Four mice per group. Data were expressed as the mean with SEM representative of the average of ∼900-1,000 MAP2 neurons comprised in 3-4 different hippocampal sections per animal. The fluorescence intensity of succinyl lysine was normalized to the number of pyramidal neurons (****: *p* < 0.0001, **: *p* < 0.01, two-way ANOVA followed by Tukey’s multiple comparisons test).
**e.** Succinylation promotes self-aggregation of tau. Tau peptides concentrations were 10 μM in presence of 2.5 μM heparin: PHF6 (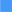), S-PHF6 (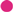), PHF6:S-PHF6 = 9:1 (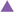), PHF6* (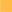), A-PHF6* (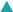). Experiments were performed in triplicate and repeated three times with similar results. All values in the present graph were expressed as mean ± SEM. All statistical analysis was implemented at time = 5,000 s (n = 3; ****: *p* < 0.0001, ***: *p* < 0.001 in comparison to S-PHF6, one-way ANOVA followed by Tukey’s multiple comparisons test).
**f-h.** Negative stain electron microscopy of *in vitro* polymerized PHFs after 24 hrs incubation. **f**: 50 μM PHF6; **g**: 50 μM S-PHF6; **h**: 50 μM mixture (PHF6:S-PHF6 = 9:1). White arrows denote paired helical filaments. Scale bar is 100 nm.
**i, j**. The width and height of the fiber helix found in polymerized PHFs after 24 hrs incubation *in vitro*. All photographed examples were measured in 3 cases, and the results averaged. Error bars represent SEM deviation from the mean (n = 3; ****: *p* < 0.0001, ***: *p* < 0.001, **: *p* < 0.01, *: *p* < 0.05, one-way ANOVA followed by Tukey’s multiple comparisons test).
**k.** Inhibition of assembly reaction of K19 and microtubules by succinylation of K19. Incubations (30 minutes) were with 30 μM succinylated K19 (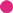) or non-succinylated K19 (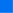). All of the experiments were performed in triplicate and repeated three times with similar results. Error bars represent SEM deviation from the mean. All statistical analysis was implemented at time = 80 min (n = 3; ****: *p* < 0.0001, one-way ANOVA followed by Tukey’s multiple comparisons test).
**l, m.** Succinylation of K19 weakens its interactions with T2R.^1^H,^15^N HSQC spectra were recorded for unmodified and succinylated K19 in the absence (black) and in the presence (red for unmodified K19, orange for succinylated K19) of T2R. **l**: Unmodified ^15^N K19 spectra (assignments for well-resolved residues as indicated) exhibit intensity loss for multiple residues including Ile^308^, Val^309^, Tyr^310^, Lys^311^ in the presence of T2R. Attenuation of resonance intensities is observed for a range of K19 resonances in the presence of T2R, and is quantified as intensity ratios (I/I_0_). **m**: Succinylated ^15^N K19 spectra exhibit less intensity loss in the presence of T2R, with residues Ile^308^, Val^309^, Tyr^310^, Lys^311^ remaining visible. Increased intensity ratios of succinylated K19 resonances in the presence of T2R compared to those for unmodified K19 indicate decreased binding upon succinylation.
**n, o:** Succinylation of K311 weakens the interactions of tau peptide (296-321) with tubulin. **n**: Comparison of 1D ^1^H spectra (black) and saturation transfer difference NMR spectra (blue) of unmodified tau peptide (296-321) in the presence of 20 μM tubulin. **o**: Comparison of 1D ^1^H spectra (black) and saturation transfer difference spectra (blue) of K311-succinylated tau peptide (296-321) in the presence of 20 μM tubulin. The tau peptide concentrations were ca. 1 mM. The signals observed in the STD spectrum of unmodified tau peptide demonstrate that it binds to tubulin. In contrast, no or weak binding was detected under these conditions for the K311 succinylated tau peptide.

In order to characterize tau succinylation in a mouse model of tangle formation, we used immunofluorescence staining to compare the presence or abeta of succinylation with that of tau oligomers (T-22)^34^ and phospho-Tau (AT8) in hippocampus from 4-month-old and 10-month-old wild type and TgP301S mice. No phosphorylated tau and few tau oligomers occurred in wild type mice (**Figure 6c, d** and **Extended Data Figure 4a, b**), but in 4-month-old TgP301S mice. Succinylation immunofluorescence signals were increased in parallel with the oligomeric tau T-22 (green) and Phospho-Tau AT8 (green) in 4-month-old TgP301S mice (**Figure 6c, d** and **Extended Data Figure 4a, b**). Thus, tau succinylation is associated with tau aggregates in TgP301S mouse model at an early stage. By contrast, a weak signal for succinylated tau occurred in 10-month-old TgP301S mice (**Figure 6c, d** and **Extended Data Figure 4a, b**), indicating a desuccinylation process may exist in the final states of tau deposition. This reflected a potential existence of succinylation-phosphorylation switch as is the case with acetylation^35, 36^.

The heparin-induced thioflavin S (ThS) tau aggregation assay was used to test the influence of tau succinylation at K311 on the ability of PHF6 to self-aggregate. PHF6* and K280-acetylated PHF6* (A-PHF6*) were also used as controls in parallel assays (**Extended Data Figure 4c**). Surprisingly, at peptide concentration of 10 μM in the presence of 2.5 μM heparin, neither PHF6* nor A-PHF6* fibrillated during an 80-min incubation period. Although PHF6* is an initiation site for tau aggregation, its potency is much lower than that of PHF6^37^, possibly explaining the observed lack of aggregation under these conditions. In contrast, PHF6 and K311-succinylated PHF6 (S-PHF6) fibrillated by 80 min and 20 min, respectively (**Figure 6e**). The aggregation of PHF6 was remarkably accelerated by the K311 succinylation. A similar enhancement of PHF6-induced aggregation occurred even with a mixture containing 90% PHF6 and only 10% S-PHF6, suggesting that succinylated tau can promote aggregation of unmodified protein (**Figure 6e**). Longer (24-hour incubations) of PHF6, S-PHF6, and a 90%/10% mixture were visualized by EM (**Figure 6f-h**). All the reactions exhibited fibrils with a typical paired helical filament appearance. However, the succinylated peptide formed abundant, short and chaotic filaments, characteristics of brain-derived Alzheimer PHFs^38–40^, while unmodified PHF6 filaments are longer and sparser, morphologies more typical of recombinant tau peptide fibers (**Figure 6i** and **6j**). Thus, both the ThS fluorescence and the EM results support an important role of succinylation in promoting pathological tau aggregation.

To understand the implications of succinylation for tau function, tubulin polymerization was assayed using the tau K19 peptide, a 99-residue 3-repeat tau microtubule-binding domain (MBD) fragment (MQ244-E372), and succinylated K19 (**Extended Data Figure 4d-f**). Native tau K19 promoted tubulin assembly as determined by increased light scattering at 350 nm, as previously reported^3, 41^, while succinyl-CoA treated K19 showed a complete suppression of tubulin assembly activity (**Figure 6k**). These findings suggest that succinylation of tau leads to a loss of normal tau function in regulating microtubule dynamics.

NMR spectroscopy was used to investigate whether succinylation mediated loss of tau microtubule assembly activity resulted from a loss of tau-tubulin interactions. The binding of the tau MBD fragment K19, to a construct, composed of two tubulin heterodimers stabilized by a stathmin-like domain (T2R), was monitored as previously described^42^. In the presence of T2R a number of NMR HSQC resonances show a reduced intensity compared to corresponding resonances of matched samples of K19 in the absence of T2R (**Figure 6i**). This decreased resonance intensity indicates an interaction between the corresponding K19 residue and the much larger T2R complex. The most highly attenuated resonances (intensity ratios < 0.2) within the MBD corresponded to residues ranging from positions 308 to 323, located in R2 of the MBD and included most of the PHF6 sequence. Succinylation of ^15^N-labeled K19 (**Extended Data Figure 4g-i**) largely abrogated intensity decreases in spectra collected in the presence vs. absence T2R, with increased intensity ratios compared to unmodified K19 across all residues (**Figure 6m**). This indicates that succinylation of K19 weakens the interaction with the T2R tubulin tetramer.

To establish whether succinylation of K311 was sufficient to specifically decrease tau-tubulin interactions, ^1^H saturation transfer difference (STD) NMR was employed to analyze the tubulin interactions of a tau peptide (residues 296-321) previously shown to comprise a high affinity microtubule binding motif within tau^43–45^. STD signals were observed for unmodified tau peptide (296-321) in the presence of tubulin (**Figure 6n**), as previously reported^45^, indicative of binding. Succinylation of residue K311 within the tau peptide (296-321) resulted in a dramatic loss of STD signals (**Figure 6o**), indicating that K311 succinylation results in a significantly decreased binding affinity of this microtubule-binding tau peptide for tubulin. The recently reported structure of tau bound to microtubules shows that K280, the R2 equivalent of K311, lies along the microtubule surface^44^. K280/K311 have their positively charged amino group in close proximity to residue E415 of α-tubulin (**Extended data Figure 4j**). Therefore, it is possible that succinylation at K311 might result in an electrostatic clash between the negatively charged succinyl group and E415 residue. A decreased affinity of K311-succinylated tau for tubulin and/or microtubules could contribute to the progression of tau pathology in AD.

## Discussion

Our study provides a system level view of the human brain succinylome in metabolic process, particularly in mitochondria, and reveals the dramatic alterations of succinylation in AD. Notably, these results demonstrate for the first time that succinylation is the key link between the signature metabolic reductions and amyloid plaques and neurofibrillary tangles in AD. The current results reveal that varied in protein succinylation, as a molecular signal, correlates with altered cerebral metabolic function in AD as the disease progresses. Other PTMs, such as ubiquitination, acetylation and phosphorylation, recently shown to affect amyloid degradation^46, 47^ and tau dysfunction^35, 46–48^, contribute to amyloidopathy and tauopathy in disease. Our findings open new areas of research on the cross talk involvon aggeregation, succinylation, acetylation, malonylation, ubiquitination and phosphorylation, which are also directly linked to metabolism and as well as implicated in amyloid and tau pathology.

The mechanisms and control of both non-enzymatic succinylation and enzymatic succinylation by cellular succinyltransferases and desuccinylases are largely unknown^49^. The data in this paper clearly demonstrate that impairing mitochondrial function decreases mitochondrial succinylation and promotes succinylation of specific non-mitochondrial proteins by altering the distribution of succinyltransferases from the mitochondria to cytosol. Precedent for this concept is provided by results showing that the movement of the DLST subunit of KGDHC to the nucleus increases histone succinylation^23^. Rotenone causes translocation of PDHC from mitochondria to other cellular compartments^50^. The decline in succinylation of mitochondrial proteins suggests that activation of descuccinylases (e.g., SIRTUINS) or general increases in NAD, a popular strategy, should be reconsidered. APP and tau were only succinylated in brains from AD patients. Thus, the modification of metabolism in disease may lead to critical succinyl-mediated modifications of extra-mitochondrial proteins including APP and tau. Preventing APP and tau succinylation and/or increasing mitochondrial succinylation may provide novel therapeutic targets for the prevention and/or treatment info of AD.

Overall, these data represent the first report of the human brain succinylome and its implications, both that for mitochondrial function as well as another for molecular pathogenesis, bot amyloidosis and tauopathy. The results provide a rich resource for functional analyses of lysine succinylation, and facilitate the dissection of metabolic networks in AD. The current studies lay the foundation for future investigation into the crosstalk between different PTMs, including acetylation, phosphorylation, and succinylation associated with AD pathology. The discovery that succinylation links mitochondrial dysfunction to amyloidosis and tauopathy may provide new molecular diagnostics as well as potential targets for therapies. Since aggregates of both succinylated Aβ and succinylated tauopathy are closely associated with β-helix dysfunction, future studies may reveal additional succinylated proteins that are associated with AD or other neurodegenerative diseases.

## Acknowledgements

The studies were supported by: NIH-NIA grants P01AG014930 (G.E.G., M.F.B.) and R37AG019391 (D.E.); R01-EY026576 and R01-EY029796 (B.T.S.); NIH SIG 1S10 OD017992-01 (S.Z.); HHSN271201300031C (V.H.); AG18877 & AG22547 (W.L.K.); Burke Neurological Institute, Weill Cornell Medicine; Integrated Medicine Research Center for Neurological Rehabilitation, College of Medicine, Jiaxing University, Jiaxing, China (Dean J. Chen).

We thank L. Cohen-Gould, MS, director of the Microscopy and Image Analysis Core Facility (Weill Cornell Medicine) for the EM and C. Bracken, PhD, director of the NMR Facility (Weill Cornell Medicine) for help with NMR experiments.

We thank E. Ivanova and Structural and Functional Imaging Core at the Burke Neurological Institute for the technical assistance.

We are grateful to the NIH Neurobiobank for providing the carefully characterized human brains.

We thank Dr. R. Kayed (Department Neurology, University of Texas Medical Branch) for kindly providing the T22 antibody for tau aggregates.

---

**Extended Data Figure 1.**
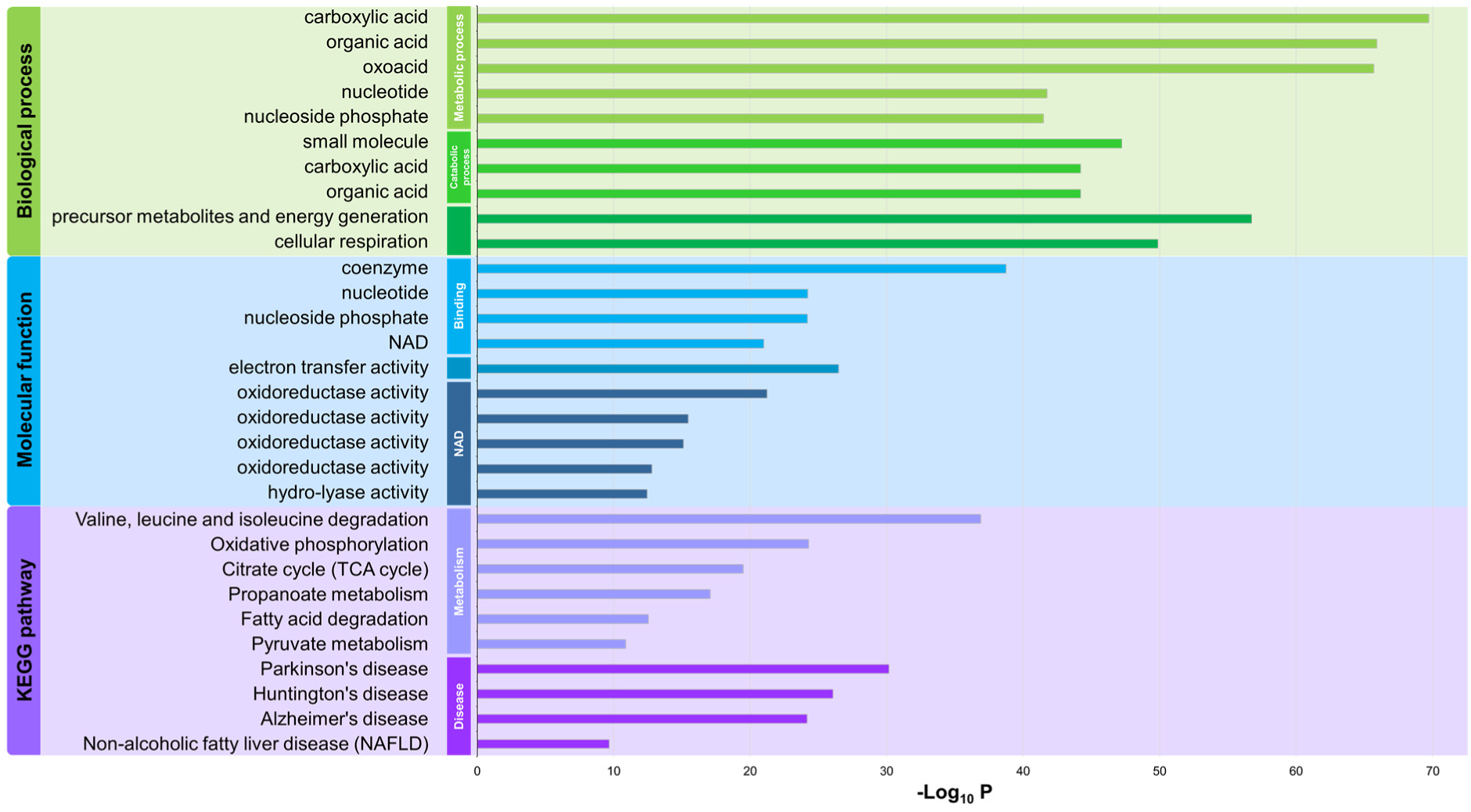
Gene ontology functional analysis of human brain succinylome. The graph shows *p*-values (step-down Bonferroni correction) for the most significant specific terms reflecting biological process (green field), molecular function (blue field) and cell component (purple field) (**Supplementary Table 3** for detail).

**Extended Data Figure 2.**
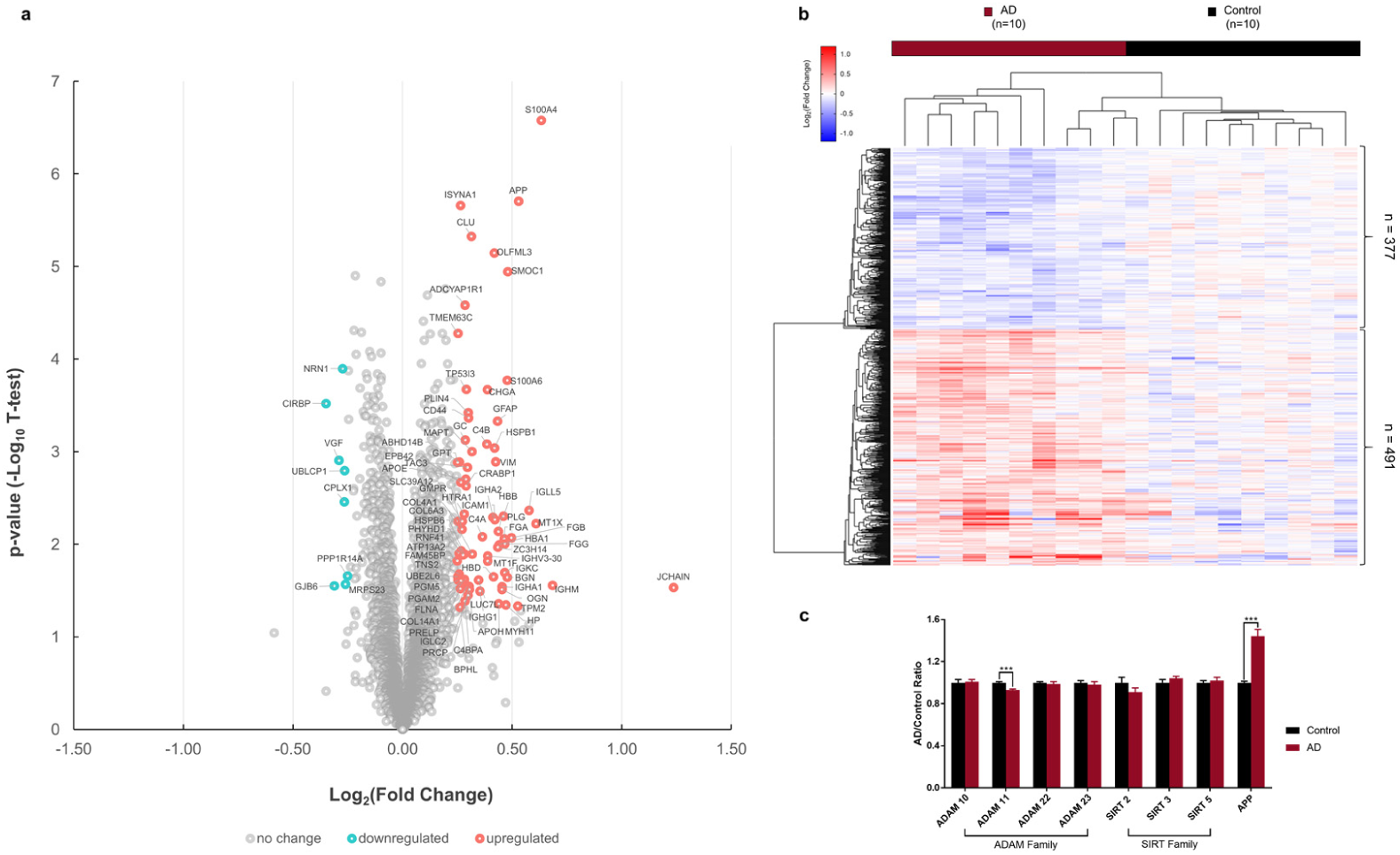
Comparison of the brain global proteomics from ten controls and ten patients with AD reveal many specific differences.

a. Volcano plot of global proteomic results comparing brains from controls and AD patients. The signal detection result shows the magnitude (mean expression difference, x-axis) and significance (− log_10_ p-value, y-axis) for brain protein level changes associations of AD. Each spot represents a specific protein. Green symbols indicate proteins that decline significantly while red symbols indicate proteins that are elevated significantly in AD brains (*p*<0.05, paired Student’s t-test, |log_2_FC|>0.25).
b. Supervised hierarchical clustering of the 868 proteins whose levels differ (*p*<0.05, two-sided Student’s t-test) between AD and control.
c. Proteomic analysis indicates that the protein levels of the α-secretase (ADAM10) are not altered in AD.

**Extended Data Figure 3.**
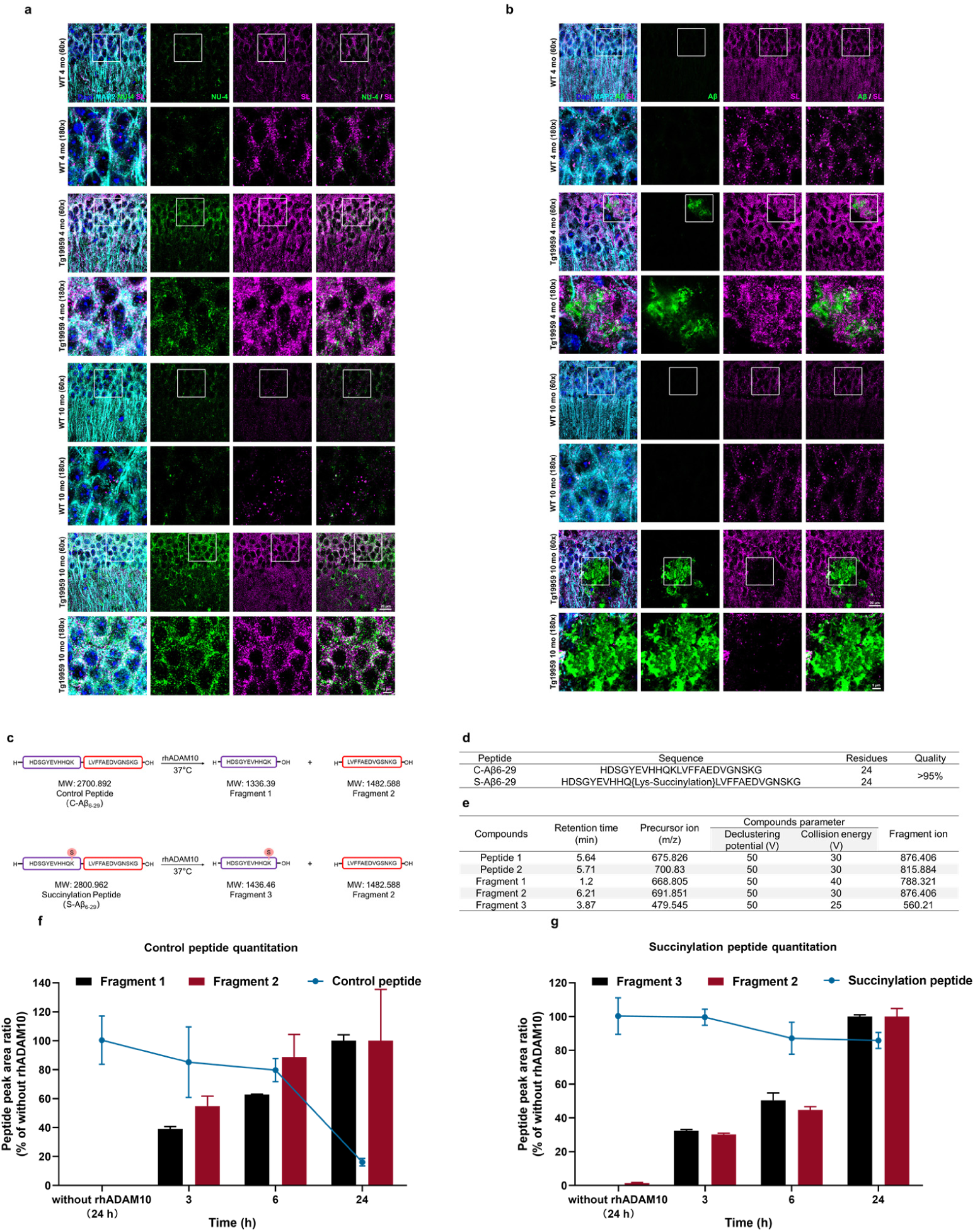
Inhibition of succinylated K687 on Aβ_6-29_ in the α-cleavage assay and succinylate Aβ_42_ using succinyl-CoA *in vitro* and its effect on ThT fluorescence assay.

a. High confocal microscope analysis results of the co-localization of succinylation and Aβ oligomers pathology in the hippocampal CA1 region sections from 4-month-old and 10-month-old Tg19959 or WT mice. NU-4 (green) staining Aβ oligomers; pan-succinyl-lysine (magenta); MAP2 (cyan); DAPi staining nuclear (dark blue).
b. High confocal microscope analysis results of the co-localization of succinylation and plaque pathology in the hippocampal CA1 region sections from 4-month-old and 10-month-old Tg19959 or WT mice. Aβ (green) staining plaque; pan-succinyl-lysine (magenta); MAP2 (cyan); DAPi staining nuclear (dark blue).
c. The schematic diagram of α-cleavage assay.
d. Properties of Aβ_6-29_ peptides used in the α-cleavage assay.
e. Multiple Reaction Monitoring (MRM) parameters with their retention time of targeted peptides and their fragments.
f. The control Aβ_42_ peptide and fragments quantitation in the α-cleavage assay. Peptide peak area ratio values were calculated and were shown relative to corresponding controls without rhADAM10. Each sample was run in triplicate. Data are expressed as the mean ± SEM.
g. The succinylated Aβ_42_ peptide and fragments quantitation in the α-cleavage assay. Peptide peak area ratio values were calculated and are shown relative to corresponding controls without rhADAM10. Each sample was run in triplicate (except for one sample from the group of succinylated peptide without rhADAM10, one sample from the group of Fragment 3 without rhADAM10, and one sample from Fragment 1 at 6 hrs were damaged) and data are means ± SEM.

**Extended Data Figure 4.**
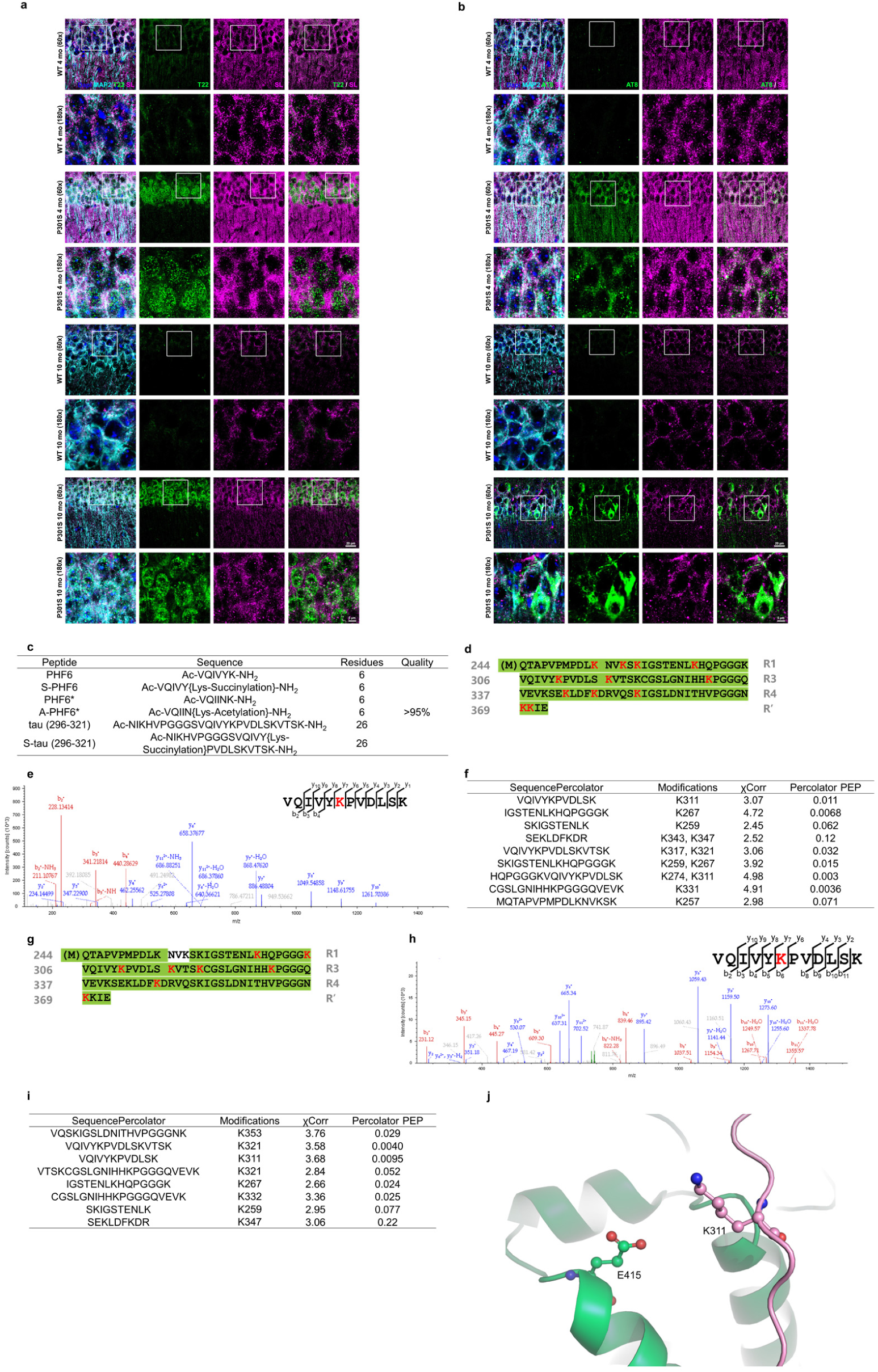
Characterization of succinylate K19 and ^15^N K19 using succinyl-CoA *in vitro*.

a. High confocal microscope analysis results of the co-localization of succinylation and tau oligomers pathology in the hippocampal CA1 region sections from 4-month-old and 10-month-old TgP301S or WT mice. T22 (green) staining tau oligomers; pan-succinyl-lysine (magenta); MAP2 (cyan); DAPi staining nuclear (dark blue).
b. High resolution confocal microscope analysis of the co-localization of succinylation and phospho-tangle pathology in the hippocampal CA1 region sections from 4-month-old and 10-month-old TgP301S or WT mice. AT8 (green) staining phospho-tau; pan-succinyl-lysine (magenta); MAP2 (cyan); DAPi staining nuclear (dark blue).
c. Properties of peptides used in the self-aggregation assay and STD NMR.
d. MS/MS identification of succ-lysines on K19 following succinylation with Succinyl-CoA *in vitro*. Residue numbering is based on the numbering of the longest tau isoform, htau40 (441 residues), and skips directly from residue 274 to 305 aa as a result of the absence of the second repeat (residues 275-305 aa). Formatting is used as follows: red, lysines (K) with succinyl group; green box, sequence covered by MS analysis.
e. Full MS and MS/MS spectra for identification and quantification of K311 succinylation on K19 following succinylation *in vitro*. b and y ions indicate peptide backbone fragment ions containing the N and C terminal, respectively. ^2+^ indicates doubly charged ions. Succ-Lysine is colored in red.
f. K19 succinylation sites identified by MS (χCorr ≥ 2.11).
g. MS/MS identification of succ-lysines on ^15^N K19 following succinylation with Succinyl-CoA *in vitro*. Residue numbering is based on the numbering of the longest tau isoform, htau40 (441 residues), and skips directly from residue 274 to 305 aa as a result of the absence of the second repeat (residues 275-305 aa). Formatting is used as follows: red, lysines (K) with succinyl group; green box, sequence covered by MS analysis.
h. Full MS and MS/MS spectra for identification and quantification of K311 succinylation on ^15^N K19 following succinylation *in vitro*. b and y ions indicate peptide backbone fragment ions containing the N and C terminal, respectively. ^2+^ indicates doubly charged ions. Succ-Lysine is colored in red.
i. ^15^N K19 succinylation sites identified by MS (χCorr ≥ 2.11).
j. Three-dimensional structure of K311 on K19 and E415 on α-tubulin during the tau-tubulin interactions.

