## Supplemental Data 1 for "Succinylation Links Metabolic Reductions to Amyloid and Tau Pathology"

### Methods

#### Human brain tissue samples

All brain tissues from Broca's area (BM-44/45, frontal lobe) were from the NIH Neurobiobank and Mount Sinai School of Medicine. All patient information including diagnosis, clinical dementia rating (CDR), age, sex, post-mortem interval (PMI) disease status and neuropathological diagnostic criteria (including mean plaques, and Braak staging) are detailed in **Supplementary Table 1**. Neuropathological evaluation of amyloid plaque distribution was based on the Consortium to Establish a Registry for Alzheimer's disease (CERAD) criteria, while the extent of spread of neurofibrillary tangle pathology was performed according to the Braak staging system.

#### Quantitative Proteomics and Succinylome

The experimental strategy and workflow are present in **Figure 1**. The analysis of two independent batches of brains tested for replicability. In each batch, samples from five patients and five controls were analyzed in parallel for TMT-based comparative proteomics and label-free quantitation of their succinylome after enrichment of succinylated peptides. After a complete analysis of the first batch, the second batch was analyzed. The mass spectrometry proteomics data have been deposited to the ProteomeXchange[1,2] Consortium via the PRIDE[3] partner repository with the dataset identifier PXD015124.

#### Protein extraction, digestion and TMT10-plex labeling

These procedures followed the PTMScan Succinyl-Lysine Motif [Succ-K] kit protocol (Cat # 13764 Cell Signaling Technology, Inc., Danvers, MA, USA). Brain tissue powders were denatured in 20 mM HEPES pH = 8.0, 9 M Urea, 1 mM Sodium orthavanadate, 2.5 mM sodium pyrophosphate, 1 mM β-glycerophate, and then homogenized with a Dounce homogenizer. After centrifugation at 20,000×g for 15 min at room temperature (r.t.), the supernatant was into a new tube. The protein concentration for each sample was determined by BCA assay using BSA as the standard.

Further processing of the proteins was then performed according to TMT Mass Tagging Kits (Thermo Fisher Scientific , Waltham, MA, USA) and Reagents protocol (http://www.piercenet.com/instructions/2162073.pdf ) with a slight modification[4,5]. A total of 50 μg protein of each sample was reduced with 10 mM DTT for 1 hrs at 34 °C, alkylated with 50 mM iodoacetamide for 30 min in the dark and then quenched with of 38 mM dithiothreitol (DTT). Each sample diluted with 50 mM tetraethylammonium bromide (TEAB) to a final concentration of 1 M Urea. Each sample was digested with 5 μg trypsin (1:10 w/w) for 18 hrs at 35 °C. Samples were then dried down in speed vac and reconstituted to a final volume of 100 μL in 50 mM TEAB prior to labeling. The Tandem Mass Tag™ (TMT™) 10-plex labels (dried powder) were reconstituted with 50 μL of anhydrous acetonitrile prior to labeling and added with 1: 2 ratio to each of the tryptic digest samples for labeling over 1 hour at r.t.. The peptides from the 10 samples (5 controls and 5 AD cases) were mixed each tag respectively with 126-tag,

127N-tag, 127C-tag, 128N-tag, 128C-tag, 129N-tag, 129C-tag, 130N-tag, 130C-tag and 131-tag. Same labeling as above was also conducted for the second sets of additional 10 samples. After checking label incorporation using Orbitrap Fusion (Thermo Fisher Scientific, San Jose, CA, USA) by mixing 5 µL aliquots from each sample and desalting with SCX ziptip (Millipore, Billerica, MA), the 10 digested samples were pooled together. The pooled peptides were evaporated to 200 µL and subjected to cleanup by solid phase extraction (SPE) on Sep-Pak Cartridges (Waters, Milford, MA). The eluted tryptic peptides were evaporated to dryness, and ready for the first dimensional LC fractionation via a high pH reverse phase chromatography as described below.

**High pH reverse phase (hpRP) fractionation**

The hpRP chromatography was carried out using a Dionex UltiMate 3000 HPLC system with the built-in micro fraction collection option in its autosampler and UV detection (Thermo Fisher Scientific, Sunnyvale, CA, USA) as reported previously[4,5]. Specifically, the TMT 10-plex tagged tryptic peptides were reconstituted in buffer A (20 mM ammonium formate pH = 9.5 in water), and loaded onto an XTerra MS C18 column (3.5 µm, 2.1×150 mm) from Waters (Waters Corporation, Milford, MA, USA) with 20 mM ammonium formate ($NH_4FA$), pH = 9.5 as buffer A and 80% ACN/20% 20 mM $NH_4FA$ as buffer B. The LC was performed using a gradient from 10-45% of buffer B in 30 min at a flow rate 200 µL/min. Forty-eight fractions were collected at 1 min intervals and pooled into a total of 10 fractions based on the UV absorbance at 214 nm and with multiple fraction concatenation strategy[4]. Each of the 10 fractions was dried and reconstituted in 125 µL of 2% ACN/0.5% FA for nanoLC-MS/MS analysis.

**Protein digestion and enrichment of succinylatedpeptides by Anti-succiK antibody beads**

For global succinylome analysis, 1 mg of proteins of each sample was reduced with 10 mM DTT for 1 hrs at 34°C, alkylated with 50 mM iodoacetamide for 1 hrs in the dark and then quenched by additional of 38 mM DTT. Each sample was diluted with 50 mM TEAB to a final concentration of 1 M Urea. Each sample was digested with 55 µg trypsin (1:18 w/w) for 18 hrs at 35 °C. The digests were cleaned up with Bond Elute C18 100 mg/mL cartridge (Agilent) and peptides were eluted with 50% ACN-0.1% TFA and dried down by a SpeedVacuum. Subsequent enrichment for succinylated peptides was conducted using a PTMScan® Succinyl-Lysine Motif [Succ-K] kit (Cat # 13764, Cell Signaling Technology, Inc., Danvers, MA, USA) following the vendor's recommended procedures. Specifically, the peptides from each sample were reconstituted in 350 µL immunoaffinity purification (IAP) buffer and transferred to the vial containing 20 µL equilibrated Succ-K motif antibody beads, incubated on a vortex mixer at 4°C for 2 hrs. After centrifugation at 2,000 g for 30 s, the beads were washed twice with 250 µL IAP buffer and three times by water. Finally, the enriched peptides were eluted three times with 55 µL of 0.15% TFA. The eluted fractions were pooled together, dried down and reconstituted in 22 µL of 0.5% formic acid (FA) for subsequent label-free quantitative analysis by nano scale LC-MS/MS.

**Nano-scale reverse phase chromatography and tandem MS (nanoLC-MS/MS)**

The nanoLC-MS/MS analysis was carried out using an Orbitrap Fusion (Thermo Fisher Scientific, San Jose, CA, USA) mass spectrometer equipped with a nanospray Flex Ion Source using high energy collision dissociation (HCD) similar to previous reports[6,7] and coupled with the UltiMate3000 RSLCnano (Dionex, Sunnyvale, CA, USA). Each reconstituted fraction (4 µL = 0.8 µg for global proteomics fractions and 20 µL for enriched SuccK samples) was injected onto a PepMap C-18 RP nano trap column (3 µm, 100 µm × 20 mm, Dionex) with nanoViper Fittings at 20 µL/min flow rate for on-line desalting and then separated on a PepMap C-18 RP nano column (3 µm, 75 µm × 25 cm), and eluted in a 120 min gradient of 5% to 35% acetonitrile (ACN) in 0.1% formic acid at 300 nL/min., followed by a 8-min ramping to 95% ACN-0.1% FA and a 9-min hold at 95% ACN-0.1% FA. The column was re-equilibrated with 2% ACN-0.1% FA for 25 min prior to the next run. The Orbitrap Fusion is operated in positive ion mode with nano spray voltage set at 1.6 kV and source temperature at 275 °C. External calibration for FT, IT and quadrupole mass analyzers was performed. The instrument was operated in data-dependent acquisition (DDA) mode using FT mass analyzer for one survey MS scan for selecting precursor ions followed by 3 second "Top Speed" data-dependent HCD-MS/MS scans in Orbitrap analyzer for precursor peptides with 2-7 charged ions above a threshold ion count of 10,000 with normalized collision energy of 38.5%. MS survey scans at a resolving power of 120,000 (fwhm at *m/z* 200), for the mass range of m/z 400-1600 with AGC = 3e5 and Max IT = 50 ms, and MS/MS scans at 60,000 resolution with AGC = 1e5, Max IT = 120 ms and with Q isolation window (m/z) at 1.6 for the mass range m/z 105-2,000 in TMT10-plex fractions.

For label-free SuccK peptides analysis, one MS survey scan was followed by 3 second "Top Speed" data-dependent CID ion trap MS/MS scans with normalized collision energy of 30%. Dynamic exclusion parameters were set at 1 within 45 s exclusion duration with ±10 *ppm* exclusion mass width. Ten samples for both control and AD cases were analyzed in Orbitrap in random order for data acquisition. All data are acquired under Xcalibur 3.0 operation software and Orbitrap Fusion Tune 2.0 (Thermo Fisher Scientific, Waltham, MA, USA).

**Data processing, protein identification and data analysis**

All MS and MS/MS raw spectra from each set of TMT10-plex experiments were processed and searched using the Sequest HT search engine within the Proteome Discoverer 2.2 (PD2.2, Thermo Fisher Scientific, Waltham, MA, USA). The *Homo sapiens* NCBI UniprotKB.fasta database containing 20,153 entries downloaded on October 17, 2016 were used for database searches. The default search settings used for 10-plex TMT quantitative processing and protein identification in PD2.2 searching software were: two mis-cleavage for full trypsin with fixed carbamidomethyl modification of cysteine, fixed 10-plex TMT modifications on lysine and N-terminal amines and variable modifcations of methionine oxidation, deamidation on asparagines/glutamine residues and protein N-terminal acetylation. The peptide mass tolerance and fragment mass tolerance values were 10 ppm and 0.02 Da, respectively. Identified peptides were filtered for maximum 1% FDR using the Percolator algorithm in PD 2.2 along with additional peptide confidence set to high. The TMT10-plex quantification method within Proteome Discoverer 2.2 software was used to calculate the reporter ion abundances that were corrected for the isotopic impurities. Both unique and razor peptides were used for quantitation. Signal-to-noise (S/N) values were used to represent the reporter ion abundance with a co-isolation threshold of 50% and an average reporter S/N threshold of 10 and above required for quantitation spectra to be used. The S/N values of peptides, which were summed from the S/N values of the PSMs, were summed to represent the abundance of the proteins. For relative ratio between the two groups, normalization on total peptide amount for each sample was applied.

For label-free quantitative data analysis of succinylated peptides, fragment ion tolerance 0.5 Da was used for the ion trap analyzer and an additional succinylation on Lys residue was specified as variable modifications. For each relative ratio of succinylated peptides/sites, no normalization was applied. The search result including ratio, *p*-value, succinylated peptide abundance for each sample was output to Microsoft Excel software for further data analysis.

**α-cleavage assay**

Synthetic C- and S-A$\beta_{6\text{-}29}$ peptides were purchased from GenScript USA Inc. (Piscataway, NJ, USA). Recombinant human ADAM10 protein (rhADAM10) was purchased from R&D Systems, Inc. (Minneapolis, MN, USA). The C-A$\beta_{6\text{-}29}$ and S-A$\beta_{6\text{-}29}$ peptides were dissolved in DMSO to reach a final concentration of 1 mM, respectively. rhADAM10 was diluted with assay buffer (25 mM Tris, 2 μM $ZnCl_2$, 0.005% (w/v) Brij-35, pH = 9.0) to reach a final concentration of 0.1 μg/μL. C-A$\beta_{6\text{-}29}$ and S-A$\beta_{6\text{-}29}$ peptide solution (10 μL) was added to 170 μL assay buffer together with 20 μL rhADAM10 or assay buffer, briefly mixed and then incubated at 37°C for different time intervals with continuous shaking. Reactions were stopped and saved at -80°C.

The assay reactions were conducted in triplicate at each time points for C-A$\beta_{6\text{-}29}$ and S-A$\beta_{6\text{-}29}$ peptides. The reaction samples were cleaned up by SPE as described previously. The eluted peptides with 50% ACN-0.1% FA were dried down and reconstituted in 100 μL of 0.1% FA with 2% acetonitrile and spiked with 0.4 pmol/μL Angiotensin I peptide (used as an Internal Standard). The samples were analyzed by LC-MS.

The resulting samples were subject to quantitative analysis for C-A$\beta_{6\text{-}29}$ peptide, S-A$\beta_{6\text{-}29}$ peptide and their respective cleaved product peptides (fragment 1, fragment 2 and fragment 3) by LC-MS/MS using an Exion LC coupled with the X500B Q-TOF mass spectrometer (SCIEX, Framingham, MA, USA). Each of 24 samples (6 μL) were injected onto a Luna $C_{18}$ column (150 × 2.1 mm i.d., 3 μm; Phenomenex) and eluted with 5% to 40% B (solvent B = 99.9% ACN-0.1% FA; solvent A = 0.1% FA) at 200 μL/min in a 6-min gradient, followed by ramp up to 70%B in 1.5min and 95%B in 0.5min. The gradient at 95%B was holding for 2 min prior back to 5%B in 0.5 min for subsequent 5.5 min of column equilibration. The X500B instrument was calibrated using the integrated calibrant delivery system (CDS). The source temperature (350°C), spray voltage (4.5 kV) and gas conditions ( with 30 arbitrary units for gas 1, gas 2 and curtain gas) were initially optimized and obtained using a positive TOF-MS survey scan by tee-in with infusion of 5 pmol/µL Angiotensin I. Quantitative data were acquired under MRM HR mode optimized for the C-A$\beta_{6-29}$ and S-A$\beta_{6-29}$ peptides using a Guided MRM-HR module. The detected MRM transitions used for quantitation with corresponding precursor ion dependent parameters were given in **Extended Data Figure 3e**. For quantitation of each peptide, the 2 most highest intensity m/z precursors ions with having 2-4 charges and 2 singly-charged product ions (whose m/z masses are larger than their precursor m/z) generated for each precursor m/z were selected for MRM HR data acquisition of all 24 samples using Sciex OS 1.3 software. The final MRM HR method includes a TOF MS scan with m/z 300 to 1,400 with 0.15 sec accumulation time followed by 12 targeted TOF-MS/MS scans for 5 peptides plus an IS peptide with 0.1 sec accumulation time and unit Q1 resolution for each scan with start/stop mass = 100/1,400 Da.

Quantitative data were processed using Analytics module in Sciex OS 1.3 software for automatically integrating peaks for each precursor m/z in TOF MS scan and MS/MS scan. One of the MRM HR transition data with highest intensity signal was used for quantitation of each peptide. A peak area ratio for each peptide against IS peptide was calculated for all 24 samples, and used for determining relative quantitation of both C-A$\beta_{6-29}$ and S-A$\beta_{6-29}$ peptides and their cleavage cleaved peptides for each time points against the time point of 24 hrs without rhADAM10 (**Figure 5e and Extended Data Figure 3f, g**).

**A$\beta_{42}$ aggregation assay**

The Beta Amyloid (1-42) Aggregation Kit (Cat # A-1170-2) was from rPeptide (Watkinsville, GA, USA). Rabbit pan-specific antisuccinyllysine (Cat # PTM-401) antibody and anti-β-Amyloid 1-16 Antibody 6E10 were purchased from PTM Biolab, Inc (Chicago, IL, USA) and BioLegend, Inc. (San Diego, CA, USA), respectively. To remove the preformed aggregates, the peptides were pretreated by the method[8] with minor modifications. A$\beta_{42}$ peptides were monomerized with a 2 hrs pre-treatment with 1, 1, 1, 3, 3, 3-Hexafluoro-2-propanol (HFIP) at 25°C, and the solvent was evaporated. This process was repeated three times to remove any preformed aggregates. Samples were stored at -20°C. Right before the experiments, 1 mg A$\beta_{42}$ was dissolved in 100 mM NaOH (1.1 mL), sonicated for 5 min and the solution was filtered through a 0.22 µm filter. The solution was diluted with Tris-buffered saline (TBS) (100 µM final concentration).

A$\beta_{42}$ (final concentration of 80 µM) was mixed with Thioflavin T (ThT, final concentration 24 µM) and succinyl-CoA (final concentration 3 mM) or ddH$_2$O in the TBS. Aliquots (100 µL) were immediately added to each well of a 96 well plate. The plate was maintained at a temperature of 37 °C for 24-48 hrs. Samples were taken during the time course of peptide aggregation to perform the experiments described below.

**A$\beta_{42}$ aggregates negative stain electron microscopy**

Eighty µM A$\beta_{42}$ peptide aggregated *in vitro* in the presence of succinyl-CoA (final concentration 3 mM) or dd$H_2O$ for 24-48 hrs, which was prepared as mentioned in the A$\beta_{42}$ aggregation assay. Samples (5 µL) were placed on 400-mesh formvar-carbon coated copper grid and allowed to settle to 2 minutes. Excess fluid was removed, and the grids were negatively stained with one drop of 1.5% uranyl acetate solution and left for 2 minutes. Then the grids were picked up and excess negative stain was wicked off, and the grids were allowed to air dry. The samples were viewed using a JEM-1400 TEM (JEOL, Ltd, Peabody, MA, USA), operated at 100 kV and imaged on a Veleta 2K × 2K CCD camera (EMSIS GmbH, Munster, Germany).

**Western blot analysis of SDS-PAGE**

A$\beta_{42}$ aggregate samples were diluted with Tricine SDS Sample Buffer (Cat # LC1676, Thermo Fisher Scientific, Waltham, MA, USA) and separated on a 10-20% Tris-Tricine gel using Tricine SDS Running Buffer (Cat # LC1675, Thermo Fisher Scientific, Waltham, MA, USA). The separated bands were transferred onto a nitrocellulose membrane and detected with the mouse monoclonal anti-Aβ oligomer specific antibody NU-2 (1:4,000; Klein's lab) and the mouse monoclonal anti-β-Amyloid antibody 6E10 (1:1,000; Cat # 803001, BioLegend, San Diego, CA, USA). The membrane was probed with s680RD Goat anti-Rabbit IgG Secondary Antibody (1:10,000; Cat # 926-68071, LI-COR Biosciences, Lincoln, NE, USA) and 800CW Goat anti-Mouse IgG Secondary Antibody (1:10,000; Cat # 926-32210, LI-COR Biosciences, Lincoln, NE, USA). The protein bands were quantified by the Image Studio Lite software (version 5.2， LI-COR Biosciences, Lincoln, NE, USA). Molecular weights were estimated using a prestained protein ladder from Bio-Rad (Hercules, CA, U.S.A.).

**Tau peptide self-aggregation assay**

Synthetic peptides were purchased from GL Biochem (Shanghai, China). In order to avoid pre-aggregation, all of these peptides were pretreated with HFIP for the aggregation assays as detailed[9,10]. The synthetic lyophilized peptides were monomerized in HFIP for 10 min. HFIP was removed by evaporation, and the peptides were dissolved in dd$H_2O$ and sonicated for 10 min. Aggregation was induced by incubating a final concentration of 10 µM peptides and 100 µM Thioflavin S at 25°C in 20 mM MOPS, 0.15 M NaCl, pH = 7.2. Heparin (2.5 µM) was added immediately prior to the readings in order to initiate the aggregation[11,12]. The data were collected in triplicate at 20 s intervals using a kinetic assay mode with a Gemini EM microplate fluorescence reader (Molecular Devices, USA). The excitation and emission wavelengths were 440 nm and 490 nm, respectively.

**Tau peptide negative stain electron microscopy**

Fifty µM synthetic peptides reassembled *in vitro* in the presence of the polynomic cofactor heparin (12.5 µM) for 24 hrs, which was prepared as mentioned in the self-aggregation assay. Samples (5 µL) were placed for 1 min on 400-mesh copper grids covered with carbon-stabilized Formvar film. Excess fluid was removed, and the grids were negatively stained with 4 successive drops of 1.5% uranyl acetate solution, blotting excess stain between drops. After the final drop and blotting, the grids were allowed to air dry. The samples were viewed using a JEM-1400 TEM (JEOL, Ltd, Peabody, MA), operated at 100 kV and imaged on a Veleta 2K × 2K CCD camera (EMSIS GmbH, Munster, Germany).

ImageJ (version 1.52a) was used for the quantification of the width and height of the fiber helix. All photographed examples were measured in 3 cases, and the results averaged. All statistical analysis and visualization were implemented in Graphpad Prism 8 (GraphPad Software, San Diego, CA, USA).

**Expression and purification of recombinant tau K19**

Recombinant K19 protein was expressed in *E. coli* BL21/DE3 cells (Novagen, San Diego, CA, USA) transfected with plasmids for the tau fragment K19 under the control of a T7 promoter, as previously described[13]. Briefly, to produce $^{15}N$-labeled protein, cells were grown in a minimal medium containing $^{15}N$-labeled ammonium sulfate as the sole source of nitrogen. Over-expression was induced with 0.5 mM IPTG at mid-log growth phase at 37°C. 3 hrs after induction, cells were collected by low speed centrifugation and lysed by sonication in a solution containing 3 mM Urea, 1 mM EDTA, 1 mM DTT, 10 mM Tris, and 1 mM PMSF followed by ultracentrifugation at 40,000 rpm for 1 hr in a Beckman ultracentrifuge using a Ti 50.2 rotor. The supernatant was dialyzed against 25 mM Tris, 20 mM NaCl, 1 mM EDTA, and 1 mM DTT before being purified by cation exchange chromatography, eluting with an NaCl gradient. Fractions containing tau K19 were pooled and dialyzed against 5% acetic acid before further purification by reverse-phase high-performance liquid chromatography on a $C_4$ column eluted with an acetonitrile gradient with 1% trifluoroacetic acid. Purified protein was dialyzed against $dH_2O$ before being lyophilized and stored at -20°C. Purity was confirmed by SDS-page.

***In vitro* succinylation of purified K19 and $^{15}N$ labeled K19**

Purified K19 (a final concentration 28 μM) was mixed with 80 mM PIPES, pH 6.8, 2 mM $MgCl_2$, 1 mM GTP, and 0.5 mM EGTA. The reaction was initiated by the addition of Succinyl-CoA (final concentration of 1.5 mM) or the equivalent amount of buffer for control. Samples were incubated for 30 min at 27°C after addition of Succinyl-CoA. The samples were concentrated by ultra-centrifugal filters for Succinyl-CoA removal, then diluted to 75 μL, and were ready for immediate use. 10 μL samples were stored at -80°C, and then digested by trypsin. After sample cleanup by SPE, the samples were analyzed by nanoLC-MS/MS analysis and database search using PD 2.2 as described above.

Purified $^{15}N$ labeled K19 (a final concentration 99 μM) was mixed with Succinyl-CoA (a final concentration 2.88 mM) or the equivalent amount of buffer for control in the same reaction buffer for 30 min incubation at 27°C. The samples were concentrated by ultra-centrifugal filters for Succinyl-CoA removal and stored on ice until use. 10 µL samples were stored at -80°C, and then digested by trypsin followed by LC-MS/MS analysis and database search using PD 2.2 as described above.

**RB3-SLD expression and purification**

Stathmin-like RB3 domain ($RB3_{SLD}$) containing two point mutations (C14A, F20W, stathmin numbering) was expressed in *E. coli* BL21/DE3 cells transfected with a PET-3d plasmid (a kind gift from Benoit Gigant) as previously described[14]. Briefly, over-expression was induced with 0.5 mM IPTG at mid-log growth phase at 37°C. 3 hrs after induction, cells were collected by low speed centrifugation and lysed by sonication in a solution containing 20 mM Tris-HCl, 1mM EGTA, 1 mM DTT at pH = 8.0, followed by ultracentrifugation for 15 min at 20,000g in a Beckman ultracentrifuge using a Ti 50.2 rotor. The supernatant was subjected to thermal denaturation (80°C for 15 min then 10 min on ice) and was centrifuged again, followed by nucleic acid precipitation using 20 mM spermine-HCL at pH = 7.0 for 30 min at 4°C with gentle agitation, and additional centrifugation (1 hr at 100,000g). The supernatant was dialyzed against 20 mM Tris-HCl, 1 mM EGTA at pH 8.0 before further purification by anion exchange, eluting with an NaCl gradient. Purity was confirmed by SDS-page before concentrating the purified protein to ca. 420 µM and buffer exchanging into 50 mM Phosphate, 0.1 M NaCl at pH = 7.0 using a gel filtration column. Final protein solution was flash-frozen in liquid nitrogen and stored at -80°C until use. SDS-PAGE analysis was used to estimate the concentration of final protein stocks.

**T2R preparation**

Stathmin-like RB3 domain (RB3) binds to heterodimeric tubulin with a 1:2 stoichiometry, forming a longitudinal dimer of tubulin dimers. For $^{1}H,^{15}N$ HSQC NMR experiments, $^{15}N$ K19, succinylated or unmodified, was dissolved to a final concentration of 48 µM in Tris-$d_{11}$ buffer (25 mM Tris-$d_{11}$, 25 mM NaCl, 2.5 mM EDTA, and 1.5 mM DTT with 10% $D_2O$ at pH 6.7) followed by addition of 53 µM of RB3-SLD stock solution. This mixture was then used to take up 4 mg of lyophilized tubulin (Cytoskeleton Inc., CO, USA) for a final T2R concentration of 50 µM.

**Microtubule assembly assay**

Tubulin (32 µM; Cat # T240-B, Cytoskeleton, Inc., Denver, CO, USA) in 80 mM PIPES, pH = 6.8, 2 mM $MgCl_2$, 1 mM GTP, and 0.5 mM EGTA was incubated for 2 min at 37°C. The polymerization was started by adding 20 µL concentrated succinylated or normal K19 (a final concentration of 60 µM). The data were collected at 350 nm (20 s intervals) using a kinetic assay mode, using a SpectraMax 250 plate reader (Molecular Devices, San Jose, CA, USA).

**NMR Spectroscopy**

Samples for NMR $^{1}H,^{15}N$ HSQC experiments of free K19 were prepared by resuspending lyophilized $^{15}N$-labeled K19 in Tris-$d_{11}$ buffer (25 mM Tris-$d_{11}$, 25 mM NaCl, 2.5 mM EDTA, and 1.5 mM DTT with 10% $D_2O$ at pH = 6.7). Samples for $^{1}H$ saturation transfer difference (STD) NMR were prepared by dissolving synthetic tau peptides (GenScript USA Inc, NJ, USA) in GPEM buffer (80 mM PIPES, 2 mM $MgCl_2$, and 0.5 mM EGTA, at pH = 6.9) to a final concentration of 1 mM. Peptide solutions were then used to take up lyophilized tubulin to a final concentration of 20 μM.

All NMR spectra were collected on a Bruker AVANCE 600-MHz spectrometer equipped with a cryogenic triple resonance probe. $^{1}H,^{15}N$ HSQC spectra were collected at 20°C with 1024 complex points in the $^{1}H$ dimension, 256 complex point in the $^{15}N$ dimension, spectral widths of 13 and 26 ppm in the proton and nitrogen dimensions, with the carrier positions on water in the $^{1}H$ dimension. Assignments were based on previously published assignments of K19[13,15]. Intensity ratios ($I/I_0$) were calculated in which $I_0$ represents the resonance intensity of residues when the protein is in a free-state, and I represents the resonance intensity of residues when the protein is in the presence of T2R (bound-state). $^{1}H$ STD data were collected at 10°C, with 4096 scans, a saturation time of 3 seconds, and on/off-resonance frequencies set to -0.5 ppm and 60 ppm, respectively.

**Bioinformatics analysis and statistical analysis**

These two coharts were combined by protein ID. Filtering base on the unique GI number or UniProtKB accession number, identical protein data in these two batch results were merged together and the final result set included only the proteins or succinylated peptides that were present in both batches.

**Subcellular localization analysis**

Subcellular localization of the identified candidates was determined using Cytoscape (version 3.6.1)[16] and stringAPP (version 1.4.0)[17] software. All the parameters were set to the default values, but only these highest compartment scores equal 5 as the high confidence localization were kept. The result was visualized in FunRich (version 3.1.3).

**Succinylated peptide sequence motif discovery and iceLogo heat map**

To determine the sequence motif for succinylation, succinylated peptides were extracted within seven amino acids upstream and downstream of identified succinylation sites. The web-based Motif-X program (http://motif-x.med.harvard.edu/)[18,19] was used to identify statistically significant motifs from the large post-translational modification peptide sequences. The motif width was chosen to be a length of 15, the occurrences number was set at 5, and the significance threshold was set at 0.00001. Since all proteins were derived from human brain, the "IPI Human Proteome" was used as the background database.

Heat map of 15 amino acid compositions is used to create an overview of all possibilities in a 2D space compared to the central succinylated site, on the IceLogo tool[20]. The precompiled Swiss-Prot composition was chosen to be Homo sapiens and the start position was set at -7. Only significantly up- and down-regulated elements, according to the given *p*-value (*p*-value = 0.05), are colored in respectively a shade of green and red. The non-regulated elements are colored black.

**Gene ontology (GO) and KEGG pathway enrichment analysis**

Analysis and visualization of Gene Ontology terms associated to succinylated proteins was performed with ClueGO (version 2.5.1)[21]. The following parameters were used when running ClueGO: Min GO Level = 3; Max GO Level = 8; Minimum Number of Genes associated to GO term = 3; Minimum Percentage of Genes associated to GO term = 4. Enrichment *p*-values were based on a two-sided hypergeometric test and Bonferroni step-down method corrected for multiple testing correction. *P*-value cut off of 0.01 and a minimum of 5 genes per ontology were used as filters prior to pruning the ontologies.

**Peptide and protein quantitation**

Perseus software[22] (version 1.6.0.7) was used for statistical analysis of the peptide and protein abundance data. In brief, quantitation was performed on these defined as quantifiable peptide and protein set including only those identified in both two batches and in a minimum of eight replicate in each group. The abundance ratio of AD and control peptides or proteins was defined as fold change (FC), and we used the logarithmic transformation of fold change ($log_2FC$) to represent the AD/control difference. Then the significant differences in succinylated peptide and protein levels were computed using two-tailed Student's t-tests and significant succinylated peptide level changes were defined as $p$-value < 0.05. Significant protein level changes were defined as $p$-value < 0.05 and $|log_2FC| > 0.25$. The search result including ratio, *p*-value, succinylated peptide abundance for each sample was output to Microsoft Excel software (version 1907) for further data analysis. All the succinylated peptides and proteins with $log_2FC$ and corresponding *p*-value are available in **Supplementary Table 4 and 6**.

**Hierarchical Clustering**

Global proteomic proteins data (rows) were clustered using uncentered pearson correlation and samples (columns) were clustered using city block distance, with an average linkage clustering method by Cluster 3.0[23]. Clustering results were visualized using Java TreeView3.0 beta01 (https://bitbucket.org/TreeView3Dev/treeview3/).

**Cell culture**

HEK293T cells (Cat # CtRL-3216) was purchased from the American Type Culture Collection (ATCC, Manassas, VA, USA) and was cultured following ATCC protocol. Unless otherwise noted, all cell culture supplies and medium were from Thermo Fisher Scientific (Waltham, MA, USA).

**Immunocytochemistry, image acquisition, and image analysis**

HEK293T cells were cultured on the Poly-D-Lysine-coated Delta T dishes (25 μg/mL) at a seeding density of $5 \times 10^4$ cells /dish. The next day, cells were treated with 100 nM Rotenone in complete medium (DMEM+10% FBS) for 60 min at 37°C. After 60 min, the cells were fixed in 4% formaldehyde in PBS (Image-iT Fixative Solution, Thermo Fisher Scientific, Waltham, MA, USA) at R.T. for 15 min. After washing in PBS and pre-incubation (1% Triton-X100 in PBS for 10 min at R.T. and PBS+2% BSA for 60 min at r.t.), they were labeled with a mixture of rabbit anti-DLST (1:600; Cat # 11954, Cell Signaling

Technology, Inc. Danvers, MA, USA) and mouse anti-CoxIV (1:1,000; Cat # 11967, Cell Signaling Technology, Inc., Danvers, MA, USA) antibodies. The primary antibodies were diluted in PBS+0.5% BSA and incubated overnight followed by a mixture of the secondary antibodies for 1 hrs at r.t.. Confocal stacks were taken using a 100× oil objective in 0.4 µm z-steps under inverted Nikon C1 confocal microscope. Image analysis was performed in Fiji software (Fiji, RRID:SCR_002285). From a confocal stack, all images (6 to 13 images) were taken for further analysis. Mitochondria were outlined using an automated thresholding of CoxIV-positive particles. Cytoplasm was defined as an area surrounding mitochondria. The created masks of mitochondria and cytoplasm were separately applied to the correspondent DLST images and integrated density of DLST was determined in mitochondria and cytoplasm, respectively. Integrated density of DLST was normalized to the area of mitochondria.

MEAN integrated intensity was calculated from the all particles in mitochondrial or cytosolic region in each field. Error bars represent SEM deviation from the mean (n = 98 fields from 19 dishes from two experiments; mitochondrial (52 control; 46 treated); cytosol (52 control; 46 control); ***: $p<0.001$, Tukey's multiple comparisons test)

**Preparation of the cytosolic and mitochondrial fractions**

HEK293T cells were cultured on the 10 cm dishes. After two days, cells (80% confluent) were treated with or without rotenone (100 nM or 5 µM) in balanced salt solution (140 mM NaCl, 5 mM KCl, 1.5 mM$MgCl_2$, 5 mM glucose, 10 mM HEPES, and 2.5 m M$CaCl_2$, pH = 7.4) for 20 min at 37°C. Cells were then washed twice with cold phosphate-buffered saline (Cat # 14190250; Thermo Fisher Scientific, Waltham, MA, USA). The cytosolic and mitochondrial fractions were prepared as described previously[24].

**Immunoprecipitation and western blots**

Samples were prepared as described[24] with the following antibodies: Pan anti-succinyllysine (1:200 for immunoprecipitation; 1:1000 for western blot; Cat # PTM-401, PTM Biolabs Inc., Chicago, IL, USA); Cox IV antibody (1:1,000; Cat # 11967, Cell Signaling Technology, Inc., Danvers, MA, USA); pyruvate dehydrogenase (PDH) antibody cocktail (1:1,000; Cat # AB110416; Abcam, Cambridge, MA, USA) which includes anti E2 (69kD), E3 bp (54KD), E1a (43.3kD), E1B (39.4 KD), and E1a; Anti-KGDHC E1k (1:1,000; Rockland, Limerick, PA, USA); Anti-KGDHC E2k (1:1,000; Rockland, Limerick, PA, USA); Anti-KGDHC E3 (Lipoamide Dehydrogenase Antibody) (1:2000; Cat # 200-4160S, Rockland, Limerick, PA, USA); β -Actin (13E5) Rabbit antibody (1:1,000; Cat # 4970, Cell Signaling Technology, Inc., Danvers, MA, USA); s680RD Goat anti-Rabbit IgG Secondary Antibody (1:10,000; Cat # 926-68071, LI-COR Biosciences, Lincoln, NE, USA) and 800CW Goat anti-Mouse IgG Secondary Antibody (1:10,000; Cat # 926-32210, LI-COR Biosciences, Lincoln, NE, USA).

All the quantification of immunoblotting analyses was taken in Image Studio Lite (version 5.2, LI-COR Biosciences, Lincoln, NE, USA), and all statistical analysis and visualizations were implemented in Graphpad Prism 8 (GraphPad Software, San Diego, CA, USA).

**Animals**

All the experiments were carried out in four and ten-month-old transgenic mouse models of AD. Tg19959 mice (that overexpress a double mutant form of the human amyloid precursor protein) were obtained from Dr. George Carlson (McLaughlin Research Institute, Great Falls, MT, USA). P301S (PS19, that overexpress the human tau gene harboring the P301S mutation) transgenic mice were purchased from The Jackson Laboratory (Bar Harbor, ME, USA). Animals were maintained under standard conditions of 12 hrs light/dark cycle, 22±1°C temperature-controlled room, and 50-70% humidity. Subjects were given *ad libitum* access to food and water. All experiments were approved by the Animal Care and Use Committee of Weill Cornell Medicine.

**Immunofluorescence**

For fluorescence analysis, tissue was washed six times in PBS for 10 min each at r.t.. After blocking in 10% normal donkey serum in PBS for 1 hrs, sections were incubated using primary antibodies (chicken MAP2, Cat # AB5543, Millipore; rabbit succynil lysine, Cat # PTM-401, PTM Biolabs; mouse NU-4 (Klein's lab); mouse Aβ, Cat #15126S, Cell Signaling; mouse T22 (Kayed's lab); mouse AT8, Cat # MN1020, Thermo Fisher Scientific, Waltham, MA, USA ) in PBS with 1% normal donkey serum overnight at 4°C. Next, hippocampal sections were rinsed three times in PBS for 10 min each and subsequently incubated with conjugated secondary antibodies (488 donkey anti-mouse, Cat # A21202, Thermo Fisher Scientific, Waltham, MA, USA; Cy3 donkey anti-rabbit, Cat # 711-165-152, Jackson ImmunoResearch; 647 donkey anti-chicken, Cat # 703-605-155, Jackson ImmunoResearch) for 2 hrs. After three washes in PBS for 10 min, the samples were mounted directly onto plus-coated slides and coverslipped using gelvatol mounting media.

**Image analysis**

The immunoreactivity of succinyl lysine was assessed in three confocal images for the CA1 region and two images for the CA2 region. For quantitative analysis, confocal fluorescence micrographs were obtained with the 40x objective lens. Representative images were captured using 60x magnification at high resolution (100 μs). Values are mean ± SEM representative of the average of ~900-1,000 MAP2 neurons or 60 Aβ plaques comprised in 3-4 different hippocampal sections per animal. The fluorescence intensity of succinyl lysine was normalized to the number of pyramidal neurons. Four mice per group. $^{****}p < 0.0001$ and $^{**}p < 0.01$ relative to WT 4 m.o. $^{****}p < 0.0001$, $^{**}p < 0.01$ and $^{*}p < 0.05$ in comparison to WT 10 m.o. $^{****}p < 0.0001$, $^{**}p < 0.01$ and $^{*}p < 0.05$ versus TG 4 m.o. (two-way ANOVA followed by Tukey's multiple comparisons test).
